# Cohort Profile: Genetic data in the German Socio-Economic Panel Innovation Sample (Gene-SOEP)

**DOI:** 10.1101/2021.11.06.467573

**Authors:** Philipp D. Koellinger, Aysu Okbay, Hyeokmoon Kweon, Annemarie Schweinert, Richard Karlsson Linnér, Jan Goebel, David Richter, Lisa Reiber, Bettina Maria Zweck, Daniel W. Belsky, Pietro Biroli, Rui Mata, Elliot M. Tucker-Drob, K. Paige Harden, Gert Wagner, Ralph Hertwig

## Abstract

The German Socio-Economic Panel (SOEP) serves a global research community by providing representative annual longitudinal data of private households in Germany. The sample provides a detailed life course perspective based on a rich collection of information about living conditions, socio-economic status, family relationships, personality, values, preferences, and health. We collected genetic data from 2,598 individuals in the SOEP Innovation Sample, yielding the first genotyped sample that is representative of the entire German population (Gene-SOEP). The Gene-SOEP sample is a longitudinal study that includes 107 full-sibling pairs, 501 parent-offspring pairs, and 152 parent-offspring trios that are overlapping with the parent-offspring pairs. We constructed a repository of 66 polygenic indices in the Gene-SOEP sample based on results from well-powered genome-wide association studies. The Gene-SOEP data provides a valuable resource to study individual differences, inequalities, life-course development, health, and interactions between genetic predispositions and environment.

## Why was this cohort set up?

Almost all human traits are partly heritable, including health outcomes, personality, and behavioral tendencies.^1,2^ All properties that make us unique as individuals are to some degree affected by random genetic variation within and between families. Moreover, genetic and environmental causes of individual differences are interrelated. For example, environmental conditions can affect how genetic differences between individuals translate into differences in socio-economic and health outcomes.^3–5^ And, genetic differences among people manifest in trait differences partly via environmental channels, for example via genetically influenced personal interests that lead to a self-selection into specific environments and reinforcement mechanisms consisting, for instance, of behaviors of parents, teachers, peers, or colleagues.^6,7^ Importantly, the fact that genetic differences are linked to differences in behavior and health does not imply simplistic biological determinism and puts no upper bound on the relevance of the environment or the possibilities for intervention.^8,9^

The heritabilities of behavioral, psychological, and economic phenotypes (e.g. educational attainment, personality, risk attitudes) and health outcomes (e.g. cardiovascular disease, dementia) are typically between 30% and 70%, with an average heritability of 49% across all traits.^2^ Thus, a substantial amount of variation in outcomes that epidemiologists and behavioral scientists study can be statistically linked to genetic differences among people. Ignoring genetics would imply that a substantial source of individual differences would remain unobserved, potentially leading to biased estimations that could prompt wrong and possibly counterproductive conclusions.^10^

Twin studies also suggest that environmental factors are important not only for social scientific outcomes, but also for a broad variety of diseases.^2^ Thus, detailed information about living conditions, attitudes, and behavior could inform health-related research questions. However, most medical research datasets only contain basic information about these factors, limiting possibilities to fully understand their importance for health outcomes.^11^

While genetically informed study designs are already common in medical research and have yielded numerous important insights into disease mechanisms,^12,13^ the use of genetic data in the social sciences is still relatively rare.^14^ Nevertheless, integrating genetic data into social-scientific research (*e.g.*, economics, psychology, sociology, political science) opens up new possibilities to (i) control for genetic confounders that are otherwise unobservable and that may lead to biased empirical results, (ii) increase the statistical power of empirical analyses by absorbing residual variance in multiple regression analyses, yielding smaller standard errors of the estimated parameters, (iii) study the interactions of genetic factors and environmental exposures, (iv) use random genetic differences among individuals to identify causal pathways, and (v) better understand how social (dis)advantages are transmitted across generations and how parents, peers, teachers, and policy makers can potentially alleviate or amplify such (dis)advantages.^14,15^ Thus, integrating genetic data into the social sciences offers researchers new tools to study questions they are interested in and to reach more robust inference on the basis of their empirical analyses.

The genetic underpinnings of behavior, socio-economic outcomes, and health are often overlapping. For example, educational attainment has substantial genetic correlations with smoking (−0.3), lung cancer (−0.4), obesity (−0.2), Alzheimer’s disease (−0.3), and longevity (+0.6),^14,16^ illustrating the complex relationships between components of genetic variation, human behavior, environmental conditions, and health outcomes.

These considerations motivated us to collect genetic data in the Innovation Sample of the German Socio-Economic Panel Study (SOEP-IS), with the goal of contributing additional value to an already existing and widely known interdisciplinary and longitudinal data set that is accessible and frequently used by the global scientific community.^17^ The addition of genetic data to this sample opens up many new research opportunities for both the medical and the social-science research community.

SOEP-IS was started in 2011 as an addition to the SOEP-Core sample, which provides representative annual data of private households in Germany since 1984.^18^ Similar to the SOEP-Core sample, SOEP-IS is a valuable data resource for researchers who want to explore long-time societal changes; relationships between early life events and later life outcomes; interdependencies between the individual and the family or household; mechanisms of intergenerational mobility and transmission; accumulation processes of resources; short- and long-term effects of institutional change and policy reforms; and migration dynamics.^18^

Besides containing a set of basic questions that are identical to the SOEP-Core, the SOEP-IS longitudinal panel survey incorporates innovative content that is purely user-designed, including measurements that go beyond the scope of standardized questionnaire formats.

As a household study, the SOEP-IS typically contains data about all household members, including a large number of mother-father-child trios, parent-offspring duos, childhood development, parenting practices, and family dynamics. Furthermore, due to the sampling method and longitudinal nature of the data, the available phenotypes in the SOEP-IS span all stages of life -- from the (pre-)natal stage, early childhood, adolescence, adulthood, all the way to retirement and the end of life (see Figure 1). We refer to the genotyped part of the SOEP-IS as the Gene-SOEP sample.

**Figure 1.**
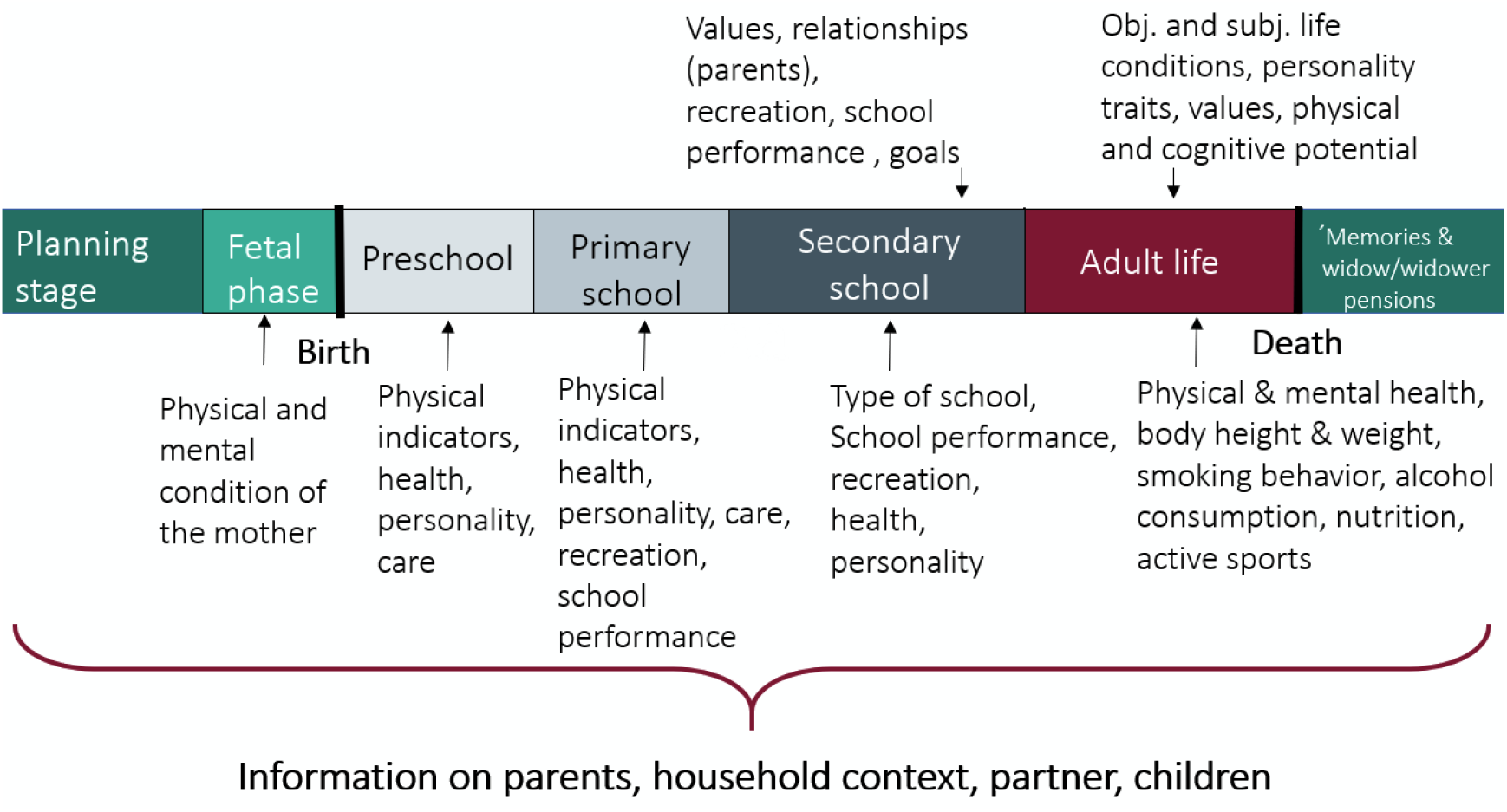
Life course perspective of the SOEP-IS sample.

Already existing genotyped cohorts in Germany (e.g. BASE-II,^19^ DHS,^20^ HNRS,^21^ KORA,^22^ SHIP^23^) focus on specific health outcomes or are limited in scope to specific regions or age groups. Thus, as of now, Gene-SOEP is the only genotyped sample that is representative of the entire German population and that contains family data as well as a rich array of longitudinal information about health, personality, family dynamics, living conditions, attitudes, and socio-economic behaviours and outcomes. This makes the sample particularly valuable to study long-term developments and the intergenerational transmission of inequalities in health and well-being. Furthermore, the sample is ideally suited to study the impact of environmental conditions that are unique to Germany, such as specific public policies and changes therein or the potential consequences of German reunification. Figure 2 shows the geographic distribution of genotyped households in the Gene-SOEP sample, illustrating the sample’s coverage of all German states and metropolitan areas (e.g. Berlin, Hamburg, Munich, Ruhrgebiet).

**Figure 2.**
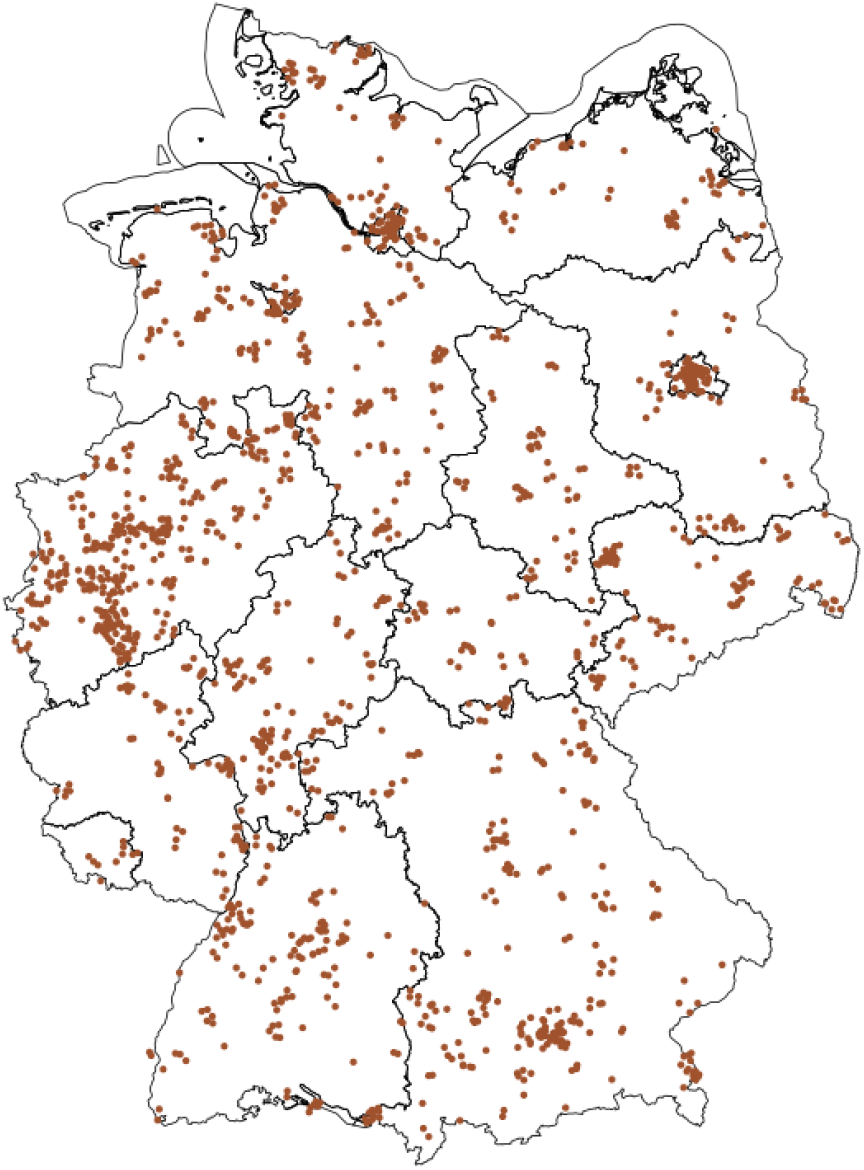
Geographic distribution of genotyped households in the Gene-SOEP sample.

To enable the collection of genetic data in the SOEP-IS, we established a research consortium of scientists from Germany (Max-Planck Institute for Human Development, German Institute of Economic Research), the Netherlands (Vrije Universiteit Amsterdam), Switzerland (University of Zurich, University of Basel), and the USA (University of Texas at Austin, Columbia University). The consortium was spearheaded by Philipp Koellinger (Vrije Universiteit Amsterdam) and Ralph Hertwig (Max-Planck Institute for Human Development). Koellinger’s team in Amsterdam developed and guided the data collection procedures, processed the collected genetic data, and generated polygenic indices for public use.

## Who is in the cohort?

The sampling and interviewing methods, as well as baseline characteristics of the sample, were previously described in detail.^17,18^ In short, SOEP-IS is based on a random sample of German households. Annual computer-assisted personal interviews are conducted face-to-face and information is collected on the household- and individual-levels (e.g. individual and household incomes). The central survey instruments are a household questionnaire. It is being answered by the household head. In addition, there is an individual questionnaire that each household member age 17 and older is supposed to answer. The surveyed information usually covers the current situation (e.g., family composition or satisfaction with life), but in some contexts it includes the past (e.g., job changes and employment biographies) and the future (e.g., expected life satisfaction in 5 years, and chance of re-employment).

The main caretaker (usually the mother) is asked about their children who are younger than 17 years. If members of an originally sampled household leave the household, (e.g. because of a divorce or children forming their own household), both the original as well as the split household are interviewed. The comprehensive tracing rules, which cover all individuals who (even temporarily) lived in SOEP households, represents a comparative advantage of SOEP compared to other household panel surveys. They allow users to track various forms of household dynamics and their implications at the household and individual level. To maintain a reasonable sample size and to address panel attrition, refreshment samples of the residential population of Germany were integrated in 2012, 2013, 2014, and 2016.

The precondition for participation in the Gene-SOEP - as part of SOEP-IS 2019 - was that the person or child lives in a participating household. 6,576 people were originally invited to participate in SOEP-IS 2019, 1,074 of whom were children. Not everyone takes part every year and there are always people who move away, die, or do not want to take part in the survey anymore. Therefore, of the original sample, 4,283 persons who were at least 17 years old (i.e., persons of survey age) as well as 875 children and youths (<17 years of age) lived in a participating household in 2019. 2,598 individuals provided a valid genetic sample, including 215 children and teenagers. A requirement for an offspring of at most 17 years of age to participate in the collection of genetic data was that both guardians agreed. The valid genetic samples were sent from the survey company Kantar Public to the Human Genomics Facility (HuGe-F) at the Erasmus Medical Center in Rotterdam for analysis.

Compared with census data (www.destatis.de), the Gene-SOEP sample is very similar to the German population in terms of age (*Mean*_census_ = 52 years vs. *Mean*_Gene-SOEP_ = 55 years), sex (51% Female_census_ vs. 54% Female_Gene-SOEP_), and living region (20% East Germany_census_ vs. 19% East Germany_Gene-SOEP_). However, residents without German citizenship are under-represented in the Gene-SOEP sample (12% census vs. 4% Gene-SOEP). Participants who agreed to donate DNA are very similar to the overall SOEP-IS sample in terms of socio-demographics, subjective health ratings, and life satisfaction (see Table 1).

**Table 1.** Descriptive statistics of the Gene-SOEP adult sample (≥ 17 years old)

|  | Total |  | Interview |  | Consent |  | Genotyped |  | Polygenic Indices Created |  |
| --- | --- | --- | --- | --- | --- | --- | --- | --- | --- | --- |
|  | Mean | SD | Mean | SD | Mean | SD | Mean | SD | Mean | SD |
| Age | 54 | 19 | 55 | 18 | 55 | 19 | 55 | 19 | 55 | 19 |
| Sex (% female) | 53 | 50 | 53 | 50 | 54 | 50 | 54 | 50 | 54 | 50 |
| East Germany (% yes) | 20 | 40 | 20 | 40 | 19 | 40 | 19 | 40 | 20 | 40 |
| German (% yes) | 95 | 22 | 96 | 20 | 96 | 19 | 96 | 19 | 98 | 16 |
| Partnered (% yes) |  |  | 41 | 49 | 40 | 49 | 40 | 49 | 41 | 49 |
| School degree: low (% yes) |  |  | 38 | 49 | 40 | 49 | 40 | 49 | 38 | 48 |
| School degree: high (% yes) |  |  | 31 | 46 | 29 | 45 | 29 | 45 | 30 | 46 |
| Employment (% yes) |  |  | 53 | 50 | 51 | 50 | 51 | 50 | 51 | 50 |
| Mean Net Income (EUR) |  |  | 1,959 | 1,304 | 1,922 | 1,300 | 1,915 | 1,258 | 1,924 | 1,263 |
| Subjective Health (1-5) |  |  | 3.33 | 0.97 | 3.34 | 0.96 | 3.34 | 0.96 | 3.33 | 0.96 |
| Life Satisfaction (0-10) |  |  | 7.54 | 1.69 | 7.57 | 1.68 | 7.59 | 1.66 | 7.58 | 1.66 |
| Observations | 5,502 |  | 4,283 |  | 2,496 |  | 2,372 |  | 2,063 |  |

Parents were somewhat hesitant to enroll their offspring (<17 years of age) for the collection of genetic data. Compared to an overall consent rate of 58% (2,496 out of 4,282 valid interviews), only 26% of the eligible offspring participated in the collection of genetic data (228 out of 875). However, offspring for whom genetic data was collected closely resemble the overall sample of offspring in the sample in terms of age, sex, geographic location, and citizenship (see Table 2).

**Table 2.** Descriptive statistics of children and adolescents (<17 years old) in the Gene-SOEP sample.

|  | Total |  | Consent |  | Genotyped Sample |  | Polygenic Indices Created |  |
| --- | --- | --- | --- | --- | --- | --- | --- | --- |
|  | Mean | SD | Mean | SD | Mean | SD | Mean | SD |
| Age | 8 | 5 | 9 | 5 | 9 | 5 | 9 | 5 |
| Sex (% female) | 49 | 50 | 50 | 50 | 50 | 50 | 50 | 50 |
| East Germany (% yes) | 18 | 39 | 19 | 40 | 18 | 39 | 20 | 40 |
| German (% yes) | 96 | 20 | 96 | 18 | 96 | 19 | 99 | 8 |
| Observations | 1,074 |  | 228 |  | 215 |  | 173 |  |

## What has been measured?

### Phenotypes

The SOEP-IS^17,24^ contains a set of core questions that are identical to about 44% of the questions asked in the SOEP-Core survey^18^, including variables such as age, gender, height, weight, education, employment status, income, life satisfaction, personality, living conditions, attitudes, preferences, and occupational classifications following the International Standard Classification of Occupations (ISCO). In addition, the SOEP-IS contains a broad range of short-term experiments and longer-term surveys that were not deemed to be suitable to the SOEP-Core survey (yet) because they pose a higher risk of refusal and panel attrition or because they deal with very specific research issues. Every year, researchers can propose new survey modules or experiments for inclusion in the SOEP-IS. The SOEP management team and the SOEP survey committee then select which modules will be included in the next survey wave.^17^ The SOEP-IS innovation modules also act as a test bed for how respondents react and some particularly important and successful modules (e.g. risk attitudes) can later be integrated into the much larger SOEP-Core survey, which collects data from ~15,000 households comprising ~26,000 individuals per year, including ~3,000 children and youths.

Health outcomes in the SOEP-IS are primarily measured based on self-reports of doctor diagnoses for a range of diseases, subjective evaluations of health and well-being, doctor visits, and the need for care. Furthermore, dried blood samples were tested for SARS-CoV-2 antibodies and oral-nasal swabs for viral RNA in a part of the SOEP-IS sample between Oct 2020 and Feb 2021, providing opportunities to study factors influencing infections with SARS-CoV-2 and long-term consequences.^25^.

Furthermore, the SOEP-IS allows users to add anonymized spatial information (e.g. federal states, spatial planning regions, counties, municipalities, and postal codes as well as GPS coordinates) and can be linked to administrative records from the German Pension Insurance and the Employer-Employee Study.^18,26^

An overview of the SOEP-IS survey content and examples of modules is provided in Box 1. The complete questionnaire of the 2019 survey wave, the 2019 SOEP annual report, and a description of all SOEP-IS modules from 2011-2018 are available online.^27–29^ An online companion for the entire data collection is available (http://companion-is.soep.de/).

#### Box 1. Summary of SOEP-IS survey content by topics and examples of modules

1. *Demography and Population* Country of origin, birth history
2. *Work and Employment* Change of job, contractual working hours, employment status, evening and weekend work, financial compensation for overtime, industry sector and occupational classification, job search, leaving a job, maternity / parental leave, registered unemployed, self-employment reasons, side jobs, supervisory position, use of professional skills, vacation entitlement, work from home, work time regulations, workload
3. *Income, Taxes, and Social Security* Asset balance, benefits and bonuses from employer, financial support received, individual gross / net income, inheritances, pension plans, social security, wage tax classification, alimony, household income and expenses, investments, repayments of loans
4. *Family and Social Networks* Circle of friends, family changes, family network, marital / partnership status, attitude toward parental role, breastfeeding, childcare, language use, leisure and activities, parenting goals, parenting style, pregnancy, relationship to other parent or child
5. *Health and Care* Alcohol consumption, health insurance, illness (self-reports of doctor diagnoses for sleep disorder, thyroid disorder, diabetes, asthma, cardiac disease, cancer, apoplectic stroke, migraine, high blood pressure, depression, dementia, joint disorder, chronic back problems, burnout, hypercholesterolemia, or other illness), reduced ability to work, sickness notifications to employer, smoking, state of health, stress and exhaustion, visits to the doctor, satisfaction with availability of care, health of child, physical and mental health of mother, nutrition, physical activity
6. *Home, Amenities, and Contributions of Private Households* Childcare hours, leisure activities and costs, school attendance by child, change in residential situation, consumption, costs of housing, home ownership / rental, loans and mortgages, birth of children, number of books in the household, persons in household in need of care, pets, residential area, size and condition of home
7. *Education and Qualification* Completed education and training, vocational training, educational aspirations for children, school enrollment of children
8. *Attitudes, Values, and Personality* Affective well-being, Big Five personality traits, depressive traits, goals in life, impulsivity and patience, income justice, life satisfaction, lottery question, optimism/pessimism, political tendency and orientation, reciprocity, religious affiliation, risk aversion in different domains, satisfaction with various aspects, social responsibility, trust and fairness, wage justice, well-being aspects, worries, temperament of child
9. *Time Use and Environmental Behavior* Time use for different activities, trip to work, use of transportation for different purposes
10. *Integration, Migration, Transnationalization* Applying for German citizenship, disadvantage / discrimination based on ethnic origins, integration indicators, language skills, native language, regional attachment, sense of home
11. *Innovative Modules* Anxiety and depression, assessment of contextualized emotions, risk attitudes, confusion, control strivings, dementia worry, determinants of ambiguity aversion, emotion regulation, expected financial market earnings, future life events, grit and entrepreneurship, happiness analyzer, impostor phenomenon, inattentional blindness, inequality attitudes, job preferences, job tasks, justice sensitivity, lottery play, multilingualism, narcissistic admiration and rivalry, ostracism, pension claims, perceived discrimination, physical attractivenes, self-control, self-evaluation and overconfidence in different life domains, sleep characteristics, smartphone usage, socio-economic effects of physical activity, status confidence and anxiety, subjective social status, work time preferences

### Genetics

DNA was extracted from saliva samples that were collected using Isohelix IS SK-1S buccal swabs with Dri-Capsules. Genotyping was carried out using Illumina Infinium Global Screening Array-24 v3.0 BeadChips, yielding raw data for 2,598 individuals and 725,831 variants, of which 688,618 were autosomal.

Call rates were smaller than 95% in 484 genotyped individuals. Further analyses revealed that the low call rates for these individuals were largely driven by interviewer effects, possibly due to not following the sample collection protocol accurately, including an incorrect use of (or entirely missing) DriCapsules that slow down the decay of DNA, low saliva and DNA yield, or polluted samples (see SI sections 2 and 3).

Since we expect that the vast majority of analyses in the genotyped SOEP-IS data will rely on polygenic indices (PGIs)^30^ rather than single genetic variant analyses, we implemented two different quality control (QC) pipelines, mild-QC and strict-QC, that are described in detail in the Supplementary Information. The mild-QC pipeline yields a higher sample size and both QC protocols yield approximately equally predictive PGIs (see below and SI section 7). Depending on the research question investigators will want to address, either the mild-QC or the strict-QC data can be used to maximize the statistical power of the analyses.

In short, both pipelines filtered out 14 individuals with sex mismatch. The strict-QC pipeline excluded 260 individuals whose genotype missingness rate was more than 20% within any chromosome and 59 individuals with excess heterozygosity/homozygosity. The mild-QC pipeline excluded only 36 individuals based on a per-chromosome missingness of more than 50% and 22 heterozygosity/homozygosity outliers. Using the mild-QC data, we identified 44 individuals of non-European ancestries, 25 of whom were available in the strict-QC sample. These individuals were also excluded from the mild- and strict-QC samples prior to imputation.

We used the Haplotype Reference Consortium reference panel (r1.1) for imputation.^31^ Imputation was completed for 2,497 individuals and 23,185,386 SNPs with imputation accuracy (*R*^2^) greater than 0.1 in the mild-QC data, and 2,299 individuals and 22,201,548 SNPs with *R*^2^>0.1 in the strict-QC data. Approximately 66% of the imputed SNPs are rare with minor allele frequencies (MAF) smaller than 0.01 and ~24% SNPs are common (MAF≥0.05; 5,463,110 in mild-QC, 5,463,110 in strict-QC). The average imputation accuracy in the mild-QC data is 0.664 and 0.695 in the strict-QC data. However, common SNPs (MAF≥0.05) are much more reliably imputed than rare SNPs, with an average imputation accuracy of 0.92 and 0.93 in the mild- and strict-QC data, respectively.

Using the imputed SNPs, we identified an additional 37 (2) individuals of non-European ancestries in the strict (mild) QC data on top of the 44 (25) individuals of non-European ancestries excluded prior to imputation, respectively. Thus, ~98% of the genotyped SOEP-IS sample is of European ancestries (see Supplementary Information section 4).

We constructed the first 20 principal components (PCs) of the genetic data for individuals with European ancestries based on ~160,000 approximately independent SNPs with imputation accuracy ≥70% and MAF≥0.01. We recommend using these genetic PCs in analyses as control variables for population stratification.^32^

### Family relationship among genotyped participants

With the exemption of parent-offspring pairs, family relationships among the participants are only surveyed via their relationship to the household head. For the genotyped participants in the SOEP-IS across the available waves from 1998 to 2019, there are 877 reported relationships for the 602 household heads. The majority (515) of these relationships are with their spouse or partner, while 346 relationships are with their child (324 biological, 11 adopted or biological, and 11 stepchild). The remaining relationships of household heads are with grandchildren (5), parents (4), a parent-in-law (1), a niece/nephew (3), a son/daughter-in-law (1), and a half sibling (1).

By using the reported relationships to the household head as well as directly reported parent-child relationships, we inferred or found 609 parent-offspring, 142 full-sibling, and 17 second-degree relative pairs in the Gene-SOEP sample. In Table S1, we compared these reported relationships to genetically inferred relationships obtained from KING^33^. We found that 19% of the pairs have inconsistencies between the reported and genetically inferred relationships. The deviations were mainly due to low genotyping quality of some individuals. When considering only the individuals whose genotyping call rate was greater than 90% using directly genotyped SNPs, 92% of the pairs in the Gene-SOEP have consistent self-reported and genetic family relationships (see section 3 and 6 of the Supplementary Information for details). We found that most of the remaining inconsistencies are due to self-reported full-siblings who are likely to be only half siblings (13 out of 97 pairs). We also found 28 self-reported parent-child pairs that appear to be non-biological from 437 pairs in total.

Furthermore, restricting to the individuals with the genotype call rate greater than 90%, we identified 88 pairs whose family relationship information was not available in the survey data. These pairs consist of 7 parent-offspring, 19 full-siblings, 33 second-degree relatives, and 29 third or fourth degree relative pairs.

Overall, out of 2,497 individuals, we genetically identified 703 individuals with at least one first-degree relatives (parent-child or full sibling) and 728 individuals that have at least one relative with at least third-degree of relatedness (first cousins or great grandparent-child). 1,769 individuals do not have close relatives on the basis of the genetic data. Note that the related pairs reported here are not mutually exclusive and some individuals can be related to multiple people.

### Polygenic indices

The effect sizes of individual single nucleotide polymorphisms (SNPs) on behavioral traits and complex diseases are usually tiny (*R*^2^ < 0.05%). Polygenic indices (PGI) aggregate the effects of observed SNPs, weighting them by their estimated effect sizes from an independent genome-wide association study (GWAS) sample.^30^ The predictive accuracy of a PGI depends on the GWAS sample size (+), the heritability of the trait (+), the number of causal genetic variants that influence the trait (-), and the extent to which the genetic architecture of the trait is similar across various environments and datasets (+).^34,35^ Thanks to rapidly growing

GWAS sample sizes in the past few years, the accuracy of PGIs has increased greatly, especially for individuals of European ancestries.^14,36^ PGIs are now beginning to capture a substantial part of the heritability of many traits, making them valuable for research in many scientific disciplines. For example, PGIs from the latest generation of GWAS analyses capture ~12% of the variation in years of schooling,^16^ ~10% of general cognitive ability,^16^ and up to 2% of various personality characteristics such as risk tolerance.^37^

This makes these PGIs useful for follow-up analyses in samples that are much smaller than the original GWAS.^14^ For example, a sample of *N =* 1,000 yields >90% statistical power to detect an association between a PGI and an outcome of interest if the PGI captures at least 1% of the phenotypic variation (two-sided *t*-test with α=0.05). An association between an outcome and a PGI with *R*^2^ = 10% can even be detected in a sample of only *N =* 110 individuals with 90% power.

We followed the methods used by Becker et al. 2021^30^ to create a repository of single- and multi-trait polygenic indices for 66 social-scientific and health traits for individuals of European ancestries in the Gene-SOEP sample. We used the largest currently available GWAS samples to create these PGIs, including publicly available GWAS summary statistics as well as non-publicly available GWAS results from 23andMe. We extended the list of 36 single-trait and 35 multi-trait PGIs in Becker at al. 2021 by including single-trait PGIs for 19 medical outcomes with well-powered GWAS summary statistics. The single-trait PGIs were based on univariate GWAS summary statistics (Table 3), whereas the multi-trait PGI were based on multivariate MTAG analyses that exploit genetic correlations between several traits to improve predictive accuracy (SI Table 3).^38^

**Table 3.** Polygenic indices in the Gene-SOEP sample from single trait GWAS results.

| Phenotype | # SNPs | GWAS *N* |
| --- | --- | --- |
| Adventurousness<sup>30,37</sup> | 1,147,160 | 557,923 |
| Age First Birth<sup>30,40</sup> | 996,620 | 169,901 |
| Age First Menses (Women)<sup>30,41</sup> | 1,142,133 | 309,043 |
| Alcohol Misuse<sup>30,42</sup> | 1,145,324 | 120,684 |
| Alzheimer's<sup>*43</sup> | 1,115,709 | 455,258 |
| Any Ischemic Stroke<sup>*43</sup> | 850,822 | 446,696 |
| Any Stroke<sup>*43</sup> | 844,962 | 446,696 |
| Atrial Fibrillation<sup>*43</sup> | 850,822 | 1,030,836 |
| Asthma<sup>30</sup> | 1,159,334 | 418,164 |
| Asthma/Eczema/Rhinitis<sup>30,44</sup> | 1,137,288 | 513,889 |
| Attention Deficit Hyperactivity Disorder (ADHD)<sup>30,45</sup> | 1,083,048 | 57,386 |
| Body Mass Index (BMI)<sup>30,46</sup> | 1,023,282 | 582,457 |
| Breast Cancer<sup>*43</sup> | 809,475 | 228,951 |
| Cannabis Use<sup>30,47,48</sup> | 1,087,000 | 156,756 |
| Cardioembolic Stroke<sup>*43</sup> | 844,996 | 446,696 |
| Childhood Reading<sup>30</sup> | 1,147,169 | 172,502 |
| Chronic Kidney Disease<sup>*43</sup> | 845,145 | 444,971 |
| Cigarettes per Day<sup>30,49</sup> | 1,150,910 | 250,057 |
| Cognitive Performance<sup>30,50</sup> | 1,148,362 | 222,914 |
| Depression*<sup>43</sup> | 835,515 | 500,199 |
| Depressive Symptoms<sup>30,51</sup> | 1,138,362 | 619,272 |
| Diastolic Blood Pressure*<sup>43</sup> | 843,500 | 757,601 |
| Drinks per Week<sup>30,49</sup> | 1,150,775 | 723,487 |
| Educational Attainment<sup>16,30</sup> | 1,147,926 | 1,047,538 |
| Ever Smoker<sup>30,49</sup> | 1,143,561 | 1,129,163 |
| Externalizing*<sup>43</sup> | 1,020,283 | 1,492,085 |
| Extraversion<sup>30,52,53</sup> | 1,113,746 | 73,906 |
| Hay Fever<sup>30</sup> | 1,159,334 | 403,179 |
| HDL Cholesterol*<sup>43</sup> | 847,159 | 187,167 |
| Height<sup>30,54</sup> | 1,022,784 | 448,198 |
| Highest Math<sup>16,30</sup> | 1,147,159 | 430,439 |
| Insomnia*<sup>43</sup> | 824,863 | 386,533 |
| Large Artery Stroke*<sup>43</sup> | 1,159,551 | 446,696 |
| Left Out of Social Activity<sup>30</sup> | 1,147,159 | 507,803 |
| Life Satisfaction: Family<sup>30</sup> | 1,159,202 | 141,864 |
| Life Satisfaction: Friends<sup>30</sup> | 1,159,184 | 138,807 |
| Longevity*<sup>43</sup> | 832,850 | 640,189 |
| Migraine<sup>30,55</sup> | 1,146,834 | 421,013 |
| Morning Person<sup>30,56</sup> | 1,123,260 | 362,840 |
| Narcissism<sup>30</sup> | 1,147,153 | 452,535 |
| Nearsightedness<sup>30,55</sup> | 1,146,729 | 301,938 |
| Neuroticism<sup>30,52,57</sup> | 1,029,577 | 389,237 |
| Number Ever Born (Women)<sup>30,40</sup> | 1,034,474 | 207,393 |
| Openness<sup>30,52,58</sup> | 987,746 | 72,308 |
| Physical Activity<sup>30,59</sup> | 1,108,549 | 140,190 |
| Religious Attendance<sup>30</sup> | 1,159,336 | 383,466 |
| Risk Tolerance<sup>30,37</sup> | 1,076,002 | 1,070,480 |
| Schizophrenia*<sup>43</sup> | 829,801 | 105,318 |
| Self-Rated Health<sup>30</sup> | 1,144,515 | 911,102 |
| Self-Rated Math Ability<sup>16,30</sup> | 1,147,159 | 564,692 |
| Small Vessel Stroke*<sup>43</sup> | 1,159,163 | 446,696 |
| Subjective Well-Being<sup>30,60</sup> | 906,574 | 502,976 |
| Systolic Blood Pressure*<sup>43</sup> | 842,552 | 745,820 |
| Triglycerides*<sup>43</sup> | 847,159 | 177,861 |
| Type 2 Diabetes*<sup>43</sup> | 851,227 | 231,426 |
| Notes: "# SNPs" is the number of SNPs that were used to construct the PGI.<br>** indicates PGIs for medical outcomes that were not originally included in Becker et al. 2021.<br>All 55 PGIs are constructed only for individuals of European ancestry ($N = 2,495$). | | |

Some of the PGIs that we created have corresponding phenotypes in the Gene-SOEP sample (e.g. educational attainment, height, BMI, risk tolerance), while others capture genetic predispositions for phenotypes that are not observable or incompletely measured (e.g. longevity, HDL cholesterol, blood pressure, and a variety of diseases including Alzheimer’s, schizophrenia, stroke, atrial fibrillation and breast cancer). These PGIs are useful proxies for unobserved traits and outcomes. For example, they can be used as control variables in studies that focus on environmental processes such as socio-economic factors that influence health^15^, to detect gene-environment interactions (e.g. heterogeneous responses to policy interventions),^5,14^ or as exogenously given proxies that do not change over the lifecourse (e.g. to study genetic predisposition for health on labor market outcomes). Finally, the availability of genetic data and PGIs from parents and their children offers exciting, new ways to disentangle genetic and environmental channels of intergenerational transmission of health, behavior, and socio-economic outcomes.^3,39^

## What has been found?

The SOEP sample is currently used by more than 9,000 registered users from 54 countries.^28^ About 300-400 publications annually are based on SOEP data, including OECD reports on the international development of inequality. Roughly 25% of these publications are in journals listed in the (social) science citation index and more than 100 publications are based on SOEP-IS data. The SOEP is also an integral database for official government reports in Germany. Major research areas that include SOEP-based publications include life course development, inequality, mobility, psychological outcomes and attitudes, migration, transition to a unified Germany, and health. Thus, the SOEP data is widely used and provides an indispensable empirical foundation to describe longitudinal developments and relationships, and a better understanding of socioeconomic processes and behavior. It is a highly valuable resource to study relationships between behavior, socioeconomic status, and health.^18^

The genetic data that we collected in the SOEP-IS sample (Gene-SOEP) is a new addition to this valuable resource. We describe first findings using the genetic data below.

### Predictive accuracy of polygenic indices for height, BMI, and educational attainment

Figure 3 shows the predictive accuracy of the PGIs for height and BMI in unrelated individuals from the Gene-SOEP sample, both for the mild and the strict version of the QC of the genetic data that we carried out. We measure the predictive accuracy of the PGIs as the difference in the explained variance (*R^2^*) before and after adding the PGI to a baseline regression that controls for a second-degree polynomial in year of birth, sex and their interactions, genotype batch indicators, and the top 20 genetic PCs. Since height and BMI were surveyed multiple times across waves, we first residualized height and BMI for age, age^2^, sex and their interactions within each wave and took the mean for each individual; then, as covariates, we used only genotype batch indicators and the top 20 genetic PCs. We obtained 95% confidence intervals by bootstrapping the sample 2,000 times.

**Figure 3.**
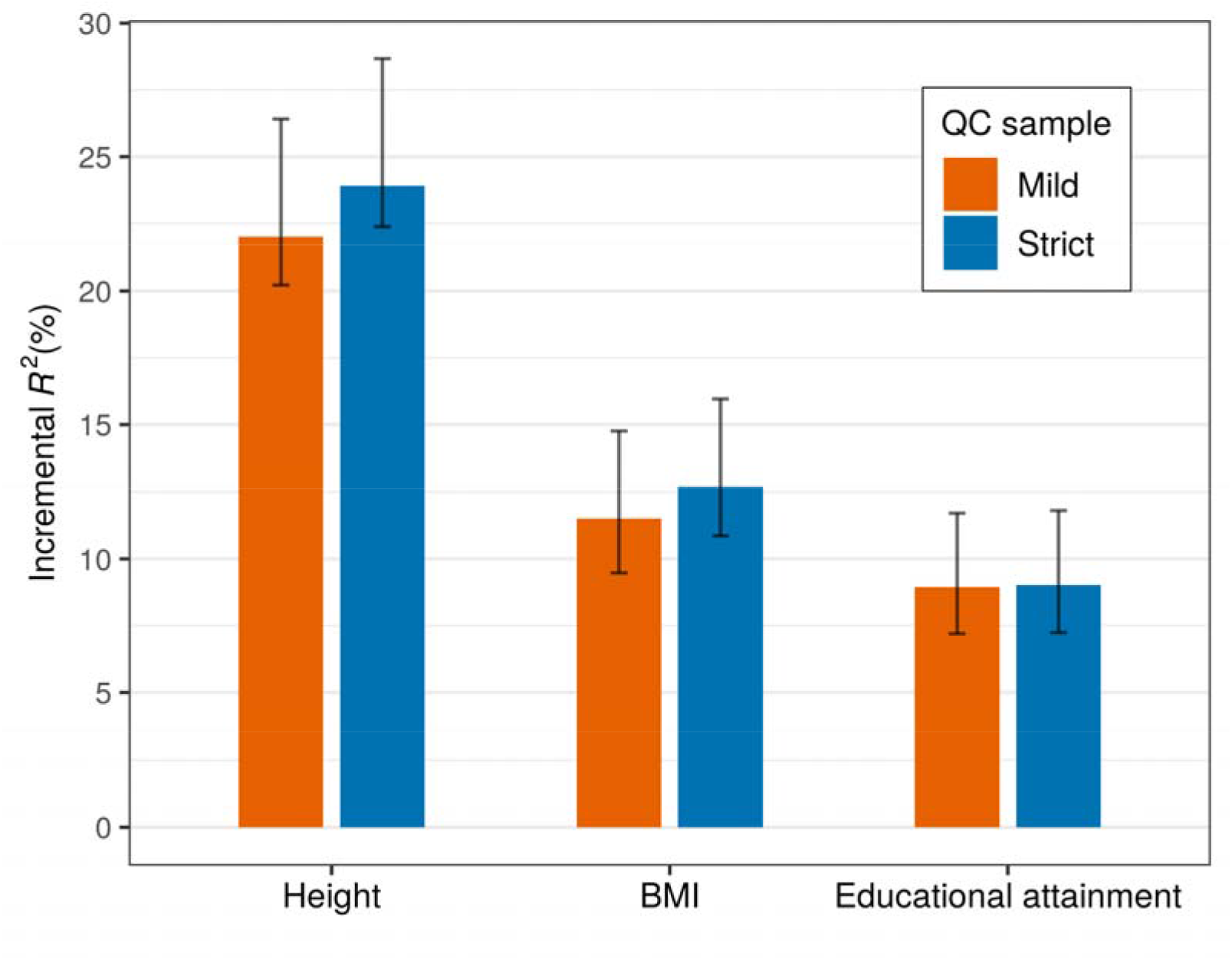
Polygenic prediction in the SOEP-IS sample. **Note**: The bars report the prediction accuracy of polygenic indices among unrelated individuals of European ancestries measured as incremental *R*^2^. The sample size of the strict (mild) QC sample is 1,904 (2,094), 1,897 (2,086), and 1,857 (2,036) for height, BMI, and educational attainment, respectively. The error bars indicate 95% bootstrapped confidence intervals with 2,000 replications.

Using this approach, the PGIs explain 22~24% of the variance in height, 12~13% of the variance in BMI, and 9% of the variance in educational attainment. Furthermore, the predictive accuracy was very similar for different levels of QC, which implies that the low genotyping quality in a part of the sample does not substantially reduce the predictive accuracy of the PGIs. Thus, researchers may choose to use the mild-QC version of the data for analyses using PGIs to take advantage of its ~10% larger sample size and the corresponding gains in statistical power.

### Genetic and environmental correlations with height and BMI

We demonstrate the advantages of combining a representative population sample with genetic data by analyzing birth year cohort trends in body height and BMI over time. Specifically, we split the Gene-SOEP sample into PGI values below and above the median for height and BMI and plotted the average residualized phenotypic values after adjusting for sex in both groups for adults >=20 years of age, binned into ten-year birth cohorts (Figures 4 and 5). Phenotypic values are residualized by regressing each observed phenotypic value on sex dummies using OLS. Each observation is assigned a residualized value which represents the remaining variation in the phenotye which cannot be predicted by sex. Residualized values are then averaged by individual across survey waves. The average residualized values for each bin are reported by the solid lines corresponding to the left axis.

**Figure 4.**
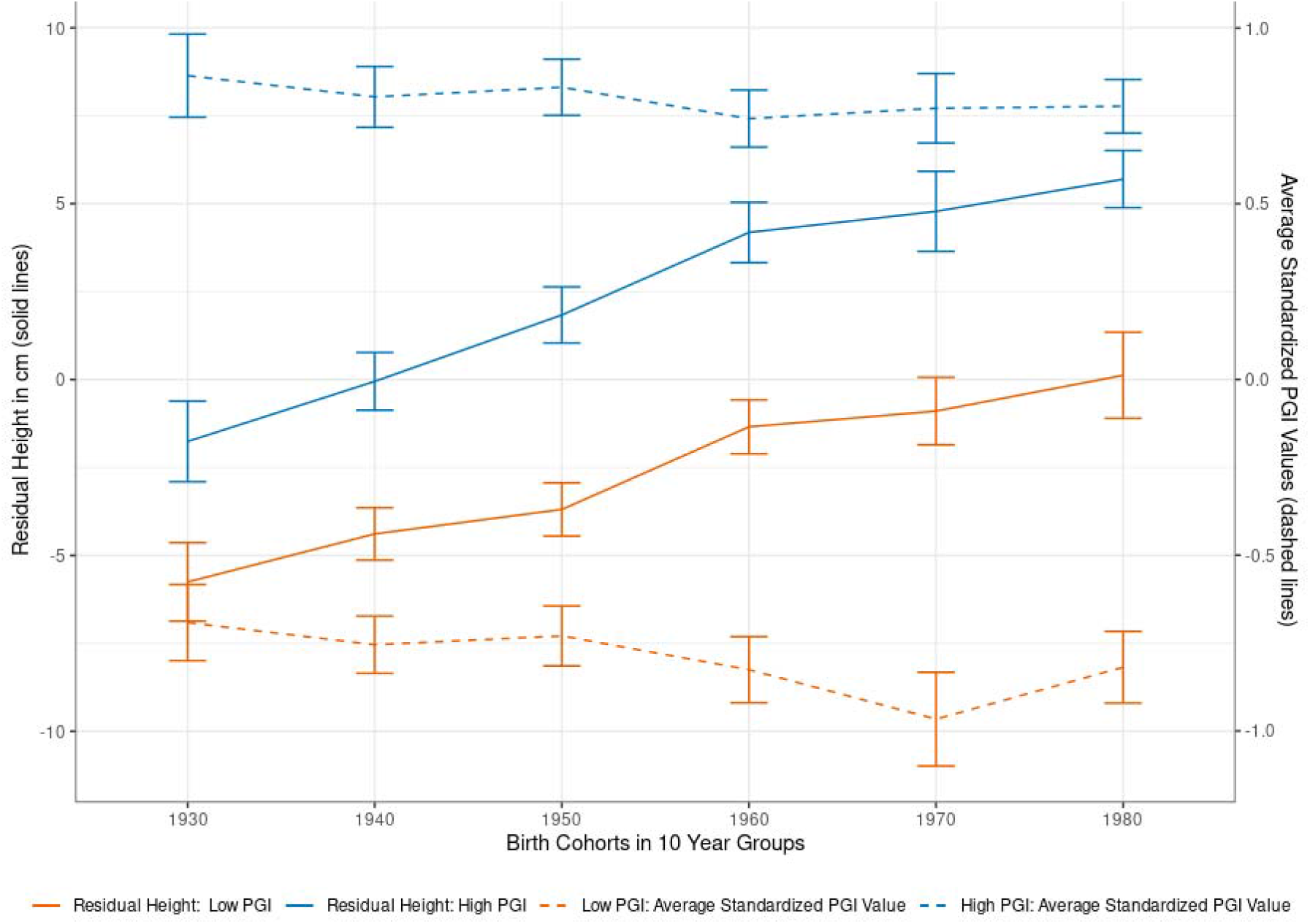
Body height by birth cohorts and PGI values. **Note**: Using the single-trait polygenic index (PGI) for body height, we split the sample of adults (older than 20 years) into two parts at the median PGI value (High PGI *N*=1,085; Low PGI: *N*=1,079). Self-reported height is residualized on sex and survey year before being averaged across survey waves. Each individual is assigned to a decadal cohort. Individuals born before between 1923 and 1939 are all in the 1930s cohort, while individuals born after 1980 are all in the 1980 group. Individuals born between 1940-1949, 1950-1959, 1960-1969, and 1970-1979 are respectively labeled as 1940s, 1950s, 1960s, and 1970s. We plotted the average observed residual height for each decadal cohort by PGI bin, along with 95% confidence intervals.

**Figure 5.**
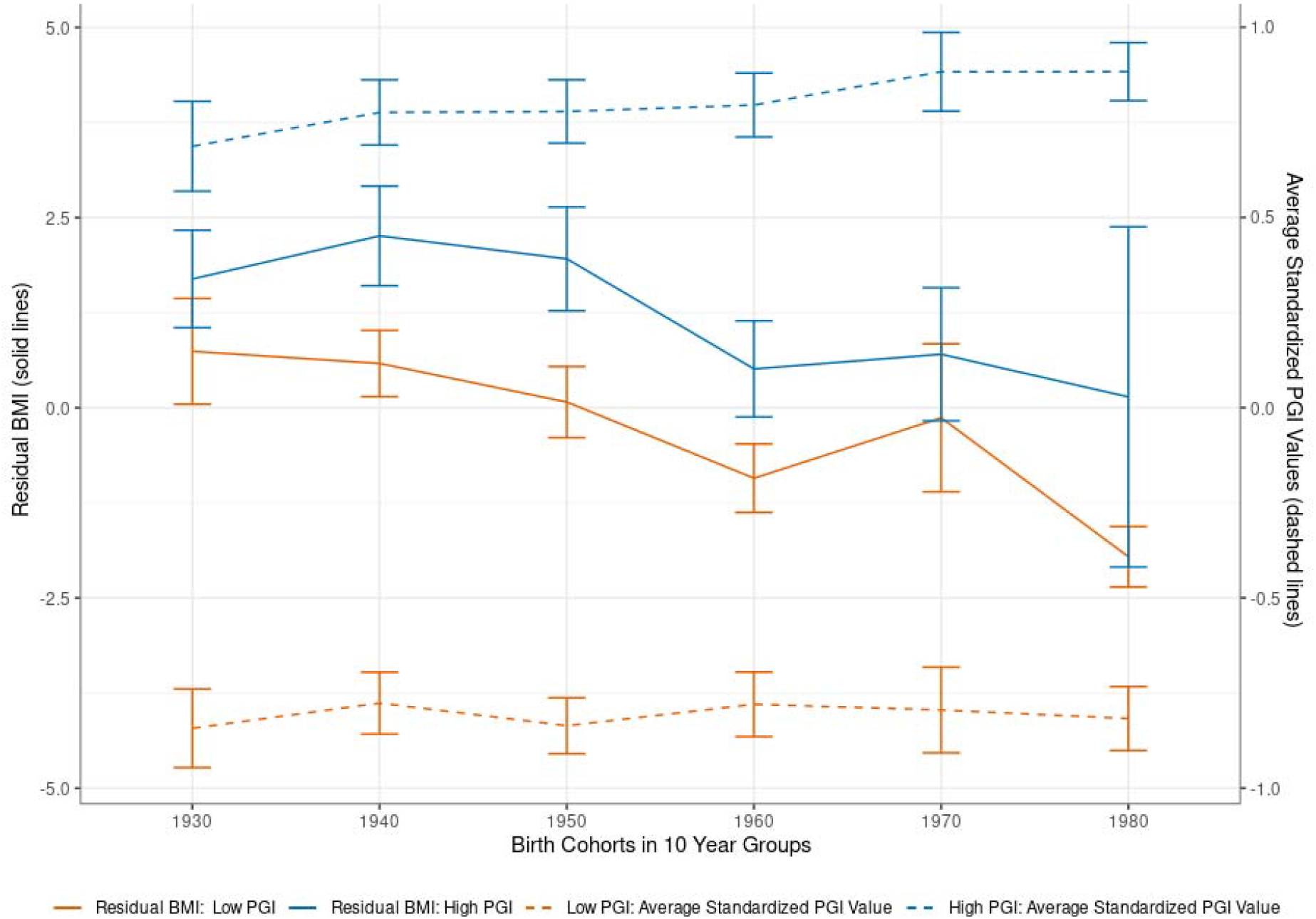
Body mass index (BMI) by birth cohort and PGI values. **Note**: Using the single-trait polygenic index (PGI) for BMI, we split the sample of adults (older than 20 years) into two parts at the median PGI value (High PGI: *N*=683; Low PGI: *N*=775). Self-reported BMI is residualized for sex and survey year before being averaged across survey waves. Each individual is assigned to a decadal cohort. Individuals born before between 1923 and 1939 are all in the 1930s cohort, while individuals born after 1980 are all in the 1980 group. Individuals born between 1940-1949, 1950-1959, 1960-1969, and 1970-1979 are respectively labeled as 1940s, 1950s, 1960s, and 1970s. We plotted the average observed residual BMI for each decadal cohort by PGI bin, along with 95% confidence intervals.

In the non-residualized data, individuals with high PGI values for height are on average 5.2 cm taller than those with low PGI height values (95% CI: 3.4 - 7.1cm). Figure 4 shows that this difference in average height by genetic predisposition is robust across birth year cohorts, reflecting a stable influence of the height PGI. Interestingly, Figure 4 also demonstrates that younger birth cohorts are on average substantially taller than older birth cohorts. For example, individuals born in the 1923-1939 birth year cohort (~84 years old on average in the 2019 survey wave) are on average 6.6 cm shorter than those born in 1980-1999 birth year cohort (~31 years old on average in the 2019 survey wave). This gain in average height of younger birth cohorts cannot be explained by observed genetic changes in the population. As Figure 4 shows through the dashed lines which correspond with the right *y*-axis, the average values of the (high and low) height PGI did not increase over time. Instead, the younger birth cohorts exhibit a slightly smaller PGI value than the older birth cohorts, possibly due to sample selection and mortality effects among older participants.^61^ In order to disentangle potential age effects from birth cohort effects, SI Table 5 presents estimates from height regressed on the standardized height PGI, birth cohort dummies, including five year age bin dummies. The results confirm a birth cohort effect on height that is separate from the genetic influences on height as well as aging effects. This implies that the substantial gains in average body height in the German population over time are partially due to improved environmental conditions, such as better nutrition and health care.^62,63^

A similar analysis for BMI (Figure 5) shows that individuals with an above-median PGI have on average also higher BMI (1.6 points higher for the High-PGI group in the non-residualized results, 95% CI 1.04 - 2.17). Both the heritability and the predictive accuracy of the PGI are lower for BMI than for height.^2,30^ Correspondingly, the average differences in BMI between the low and the high PGI group are not statistically significant for all birth year cohorts. Yet, similar to the analyses on height, we also observe birth cohort effects on BMI that cannot be explained by observed genetic variation in the BMI PGI. Individuals born in the youngest birth cohort (1980-1999, ~31 years old) have an average BMI that is 2.3 points lower than those in the oldest birth cohort (1923-1939, ~84 years old). The higher BMI in the older birth cohorts is not due to observed genetic changes in the population over time. In fact, the average PGI is slightly lower in the older birth cohorts than in the younger ones, again possibly due to sample selection and mortality effects among older participants.^61^ SI Table 6 presents regression results from a robustness check that also included 5-year age bins as control variables, again confirming birth cohort effects that cannot be explained alone by aging or observed genetic variation. Thus, the higher BMI in the older birth-cohorts is likely to be caused by a combination of environmental effects such as differences in living conditions, socio-economic effects,^64^ or nutrition.^65^

The broad set of PGIs we created are a valuable resource for research on inequalities in socio-economic and health outcomes. Previous research has demonstrated that the genetic architectures of socio-economic, behavioral and health outcomes are often substantially overlapping.^14,66,67^ This implies that PGIs for socio-economic or behavioral traits can also be proxies for health outcomes.

This is demonstrated in Figure 6, which presents the effect size from regressions of self-rated health on 28 single-trait PGIs (out of 55 tested single-trait PGIs overall) whose estimated standardized coefficients are greater than |±0.1| All regressions controlled for five year age bins, sex, and their interactions, and the first 20 genetic principal components. 18 PGIs are statistically distinguishable from zero after a Bonferonni correction for 55 tested hypotheses (marked with *).

**Figure 6.**
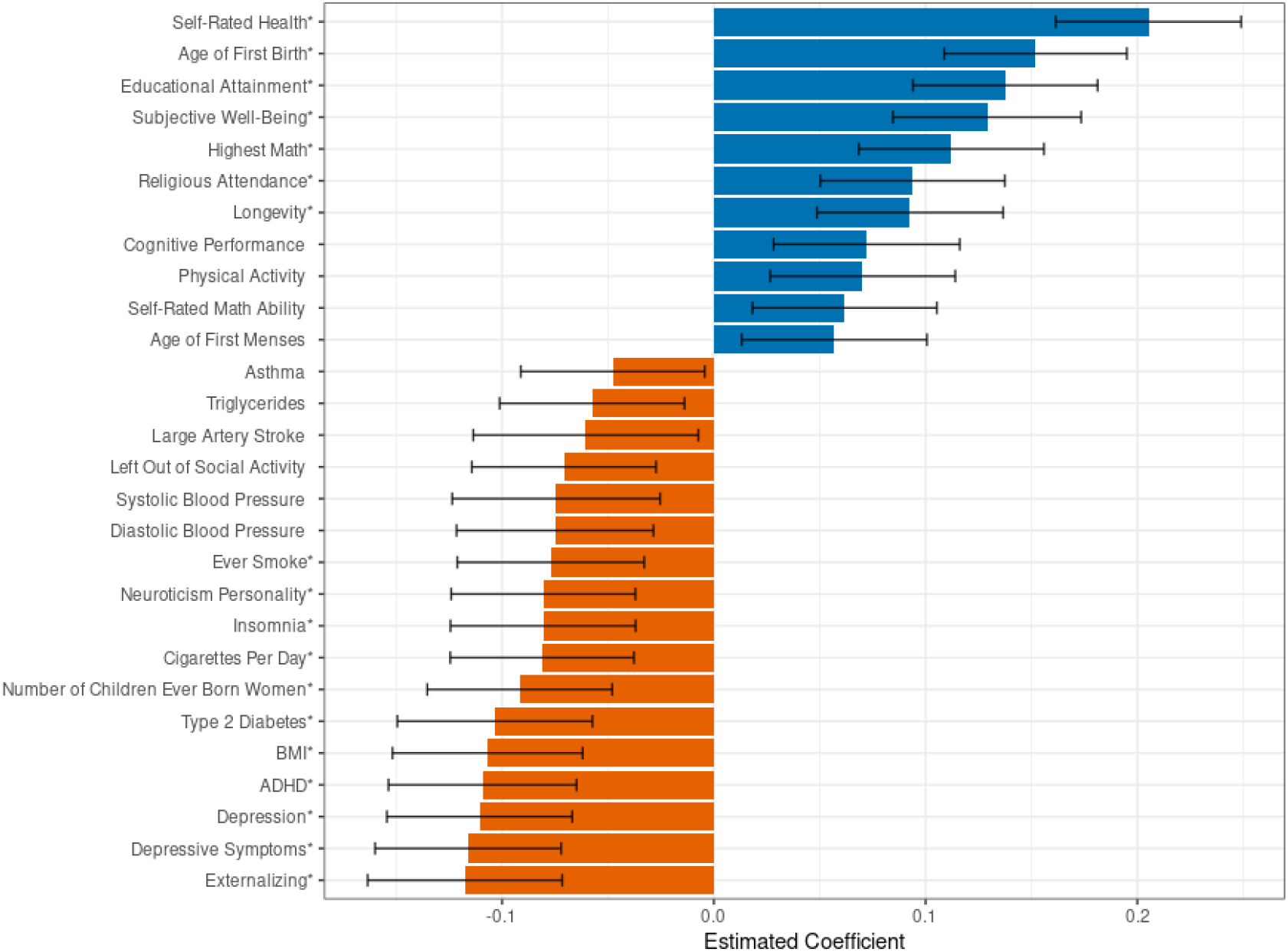
Associations between polygenic indices and self-rated health. **Note**: Analyses in the Gene-SOEP sample, *N* = 2,060. Self-rated health is measured by a 5-point Likert scale where a 1 indicates poor health and a 5 indicates very good health. Each self-rated health observation is regressed on five year age-bin dummies, sex dummies, and the interaction of sex and age bin dummies. We take the estimated residual from the previous regression, compute the average residual value for each individual, and regress each PGI along with 20 genetic principal components on these residuals where each individual has one observation. The estimated standardized betas from each PGI are reported in the figure. The figure represents 28 single-trait PGIs with an effect size of greater than |±0.1|, out of 55 single-trait PGIs overall. PGIs marked with an * are statistically distinguishable from zero after a Bonferonni correction. Error bars represent a 95% confidence interval around the estimated beta for each PGI.

We find positive associations between self-rated health and PGIs for self-rated health, age at first birth, educational attainment, subjective well-being, highest math class taken, religious attendance, longevity, cognitive performance, physical activity, self-rated math ability, and age at first menses. Furthermore, we find negative health correlations of the PGIs for externalizing, depression, ADHD, number of children ever born, insomnia, neuroticism, smoking, and being left out of social activities - all of which are PGIs for behavioral, social, or cognitive phenotypes. Moreover, the PGIs for BMI, high blood pressure, type 2 diabetes, large artery stroke, triglycerides and asthma all have the expected negative correlations with self-rated health.

## What are the main strengths and weaknesses?

Major strengths of the Gene-SOEP data include:

i. the sample selection, which yields the only currently genotyped sample that is representative of the entire German population;
ii. the longitudinal nature of the data with annual observations since 2011 (for a subset of individuals and phenotypes, annual observations even go back to 1998);
iii. the rich questionnaire content, including self-reported health outcomes and detailed information on socio-economic status, living conditions, family dynamics, personality, preferences and attitudes is another major strength of the data;
iv. the possibility to use detailed geo-coding, standardized occupation codes, and links to external databases such as the German Pension Insurance and the Employer-Employee Study;
v. the broad set of state-of-the-art polygenic indices that we created, which lower the entry barriers for researchers to use genetically informed study designs;
vi. the continuing annual collection of data that also allows researchers to integrate new survey modules, biomarkers, and experiments in the future by following the application procedures of the SOEP-IS management team;^17^
vii. the household sampling procedure that collects data on all family members. The Gene-SOEP sample contains 501 parent-offspring pairs, 152 parent-offspring trios, 107 full-siblings, and 12 second degree relatives (including half-siblings) with matching self-reported and genetically-inferred relationships. This data structure enables genetically informed studies on a wide range of research topics, including the intergenerational transmission of inequalities in health and well-being as well as studies that identify how environmental factors such as parenting style influence the developmental trajectory of children and youths;
viii. the availability of epigenetic data, which will be added for a substantial part of the Gene-SOEP sample in the near future, further increasing research opportunities on the relationships between social environment and physical health;
ix. the possibility to extend the collection of genetic data to all SOEP surveys, which would substantially increase the available sample size for genetically informed analyses.

Compared to other datasets that were included in the Polygenic Index (PGI) Repository of the SSGAC,^30^ the Gene-SOEP is the only German sample and it has the broadest coverage of social scientific outcomes, many of which have been repeatedly collected over time. Although the sample size of the Gene-SOEP is larger than several other studies included in the PGI Repository (e.g. Dunedin, E-Risk, Texas Twins), we still caution that researchers using the data should pay attention to statistical power in their analyses. In particular, the sample size may be too limited for analyses of single genetic variants or sub-parts of the sample (e.g. specific age groups or geographic areas). A further limitation is that a part of the sample (19%) did not pass the strict quality control thresholds of genetic data that are usually employed in genetic epidemiology (call rates > 95%). However, our mild-QC pipeline still enables the use of well-performing PGIs in 2,495 individuals (96% of the successfully genotyped sample).

Another possible limitation is that the currently available health outcomes are limited in detail and based on self-reports rather than detailed digital health records. Future expansions of the collected health data would further increase the utility of the SOEP samples for epidemiological research.

## How can I access the data?

The collected phenotypes from all SOEP samples can be accessed via user agreements with DIW Berlin (https://www.diw.de/en/diw_01.c.601584.en/data_access.html). The raw genetic data from Gene-SOEP will be stored on the European Genome-Phenome Archive (https://ega-archive.org/) from 2023 onwards and data access applications will be handled by DIW Berlin. Raw genetic data will need to be stored on high-security servers that meet the technical and organizational security measures required by the General Data Protection Regulation of the European Union. From 2023 onwards, DIW Berlin will also share the genetic PCs and all PGIs that were constructed in a standard phenotype file (e.g. in Stata, SPSS, or CSV formats). This version of the data also includes an indicator for individuals that did not pass the strict-QC pipeline. This allows users to decide whether they prefer to conduct their analyses using the full sample for which PGIs were constructed or the slightly smaller set that passed strict QC. DIW Berlin will also share family relationship data for each related pair, inferred from both the survey and genetic data, which will also contain genetic kinship estimates.

## Funding

The data collection was supported through the German Research Foundation (Leibniz Prize to Ralph Hertwig), a European Research Council Consolidator Grant (647648 EdGe to Philipp Koellinger), the Jacobs Foundation (EMTD, KPH, DWB), NIH/NICHD grant R01HD092548 (K. Paige Harden), the NORFACE DIAL Grant 462-16-100 (Pietro Biroli), the University of Basel (Rui Mata), and the Canadian Institute for Advanced Research (Daniel W. Belsky). The Population Research Center at the University of Texas at Austin is funded by NIH center grant P2CHD04284.

## Acknowledgements

We are deeply indebted to all individuals who have agreed to participate in the German Socio-Economic Panel Innovation Survey. The study received ethical approval by the Research Ethics Review Board of Vrije Universiteit Amsterdam, School of Business and Economics (application number 20181018.1.pkr730) and the Ethics Council of the Max Planck Society (application number 2019_16). The construction of polygenic indices in this study were made possible by the generous public sharing of summary statistics from published GWAS from many research consortia. We would like to thank the studies that made these consortia possible, the researchers involved, and the participants in those studies, without whom this effort would not be possible. We would also like to thank the research participants and employees of 23andMe for making this work possible.

## Conflicts of interest

None declared.

## Key messages

⍰ Genetic data has been successfully collected for 2,598 participants of the German Socio-Economic Panel Innovation Survey wave 2019-2020. The genotyped part of the sample (Gene-SOEP) contains 501 parent-offspring pairs, 152 parent-offspring trios, 107 full-siblings, and 12 second degree relatives (including half-siblings) with matching self-reported and genetically-inferred relationships. These family relationships are partially overlapping, e.g. the 152 parent-offspring trios are included in the 501 parent-offspring pairs.
⍰ The Gene-SOEP is currently the only genotyped sample that represents the entire German population.
⍰ Annual surveys are conducted since 2011 for all household members, generating a rich and detailed portrait of the past and current living conditions of the sample participants, including socio-economic status, well-being, health, personality, economic preferences, opinions, family dynamics, and child development.
⍰ We created a repository of 66 polygenic indices (PGIs) for social-scientific and health traits in the Gene-SOEP sample based on results from well-powered genome-wide association studies. This repository provides a valuable resource for interdisciplinary research in the medical and social sciences.
⍰ Using PGIs for body height and BMI, we demonstrate both genetic and environmental influences on the distribution of these phenotypes across different birth cohorts.

## Notes

### Competing Interest Statement

The authors have declared no competing interest.

