## Supplementary Information for "Cohort Profile: Genetic data in the German Socio-Economic Panel Innovation Sample (Gene-SOEP)"

### 1. Genotyping

We collected DNA in the SOEP Innovation Sample[1] during the 2019 annual household survey. The DNA data collection project („Wie uns die Gene prägen“) was approved by the SOEP management team as a user-proposed innovation module. The project received ethical approval by the Research Ethics Review Board of Vrije Universiteit Amsterdam, School of Business and Economics (application number 20181018.1.pkr730) and the Ethics Council of the Max Planck Society (application number 2019_16). Data collection took place between September 2019 and March 2020. Participants were visited by a trained Kantar GmbH interviewer who asked survey questions and collected saliva samples. Participation in the DNA collection was completely voluntary. Each household received information about the goals, methods, risks and benefits of the DNA collection via mail prior to the household visit, as well as at the beginning of the interview. Household members who chose to participate in the DNA collection signed a consent form and a data protection declaration according to article 7 of the EU General Data Protection Regulation. Parents had the option to enroll and provide consent for their under-aged children.

Saliva samples were collected at the end of the interview using Isohelix IS SK-1S buccal swabs. To avoid untimely decay of DNA, the buccal swabs were sealed with Isohelix Dri-Capsules in 5mm tubes. Samples were sent via mail to the Kantar GmbH office in Munich and were forwarded from there to the Human Genomics Facility (HuGe-F) at the Medical Center of Erasmus University Rotterdam. HuGe-F is a certified laboratory that operates under a Propel certificate by Illumina and works according to standard operating procedures. ICT/data handling was performed following ISO certificates ISO/IEC 27001:2013 and NEN 7510:2011. HuGe-F extracted DNA and genotyped samples using Illumina Infinium Global Screening Array-24 v3.0 BeadChips[2] between April and July 2020. The laboratory also performed a first round of data quality control and a preliminary imputation of the genetic data using the 1000 Genomes P3v5 reference panel.

DNA extraction was successfully completed for 2,598 individuals and 725,831 genetic variants (including SNPs and indels) were called by the laboratory. The quality control pipeline of the laboratory identified 2,545 individuals with European ancestries and 544 pairs with first-degree relatedness.

### 2. Genotyped data description

Figure S1 shows the minor allele frequency (MAF) distribution of the genotyped variants. 13.5% of the variants are rare variants (MAF < 1%), 35.4% low-frequency variants

(1%<MAF≤5%), and 51% are common variants (MAF>5%). The majority of the variants were called with at least 90% call rate (Figure S2A). Yet, genotype missingness rates were problematically high for almost 20% of the sample (Figure S2B). 502 individuals had a genotype missingness rate higher than 10%, considering only autosomal biallelic single nucleotide polymorphisms (SNP) with MAF > 1%.

We first investigated whether the low call rates are more prevalent in a specific MAF range (Figure S3). Although the mean call rate was lower around MAF=10%, the differences across the MAF spectrum were not large. Instead, the low call rates appeared to be due to issues during the saliva collection. Indeed, the laboratory reported a low DNA yield (<300 ng) for 301 individuals and a low DNA concentration (<3 ng/μL) for 273 individuals. As demonstrated by Figure S4, we found that some interviewers may have failed to follow the saliva collection procedure with sufficient care. 19% of the variation in the sample call rates could be explained by the variation between different interviewers. A follow-up investigation revealed that some interviewers did not properly seal the buccal swabs or forgot to include the Dri-Capsule, which may have led to pretimely decay of the collected DNA and in turn, low sample call rates.

### 3. Quality control and imputation

The laboratory implemented their standard quality control (QC) protocol, which was developed primarily for genetic discovery studies on medical phenotypes. The protocol drops variants with a call rate lower than 95% and Hardy-Weinberg equilibrium exact test *P* value lower than $10^{-4}$ as well as removing samples whose genotype missingness rate was higher than 5%. This procedure led to a large reduction in sample size, leaving only 2,114 individuals with 590,377 variants (including SNPs and indels). The large reduction in sample size led us to develop a different protocol tailored to handle the above issues with the saliva collection in SOEP-IS, described next.

We expect the most common use of the genetic data in SOEP-IS to be analyses using polygenic indices (PGIs). Therefore, we concluded that the strict standard QC protocol of the laboratory, albeit a suitable one for analyses of single genetic variants, may not be strictly necessary for analyses that utilize PGIs. PGIs aggregate the effects of hundreds of thousands of genetic variants. Thus, we expected that the accuracy of PGIs is more robust to low genotyping call rates and the drop in imputation accuracy resulting from it than analyses of single genetic variants. In order to assess whether this is indeed the case, we decided to apply two different sets of sample-level quality control filters (henceforth referred to as mild and strict QC) for all 2,598 individuals and 688,618 autosomal variants (including indels). Based on the mild and strict QC results, we also performed separate imputations. With these two sets of imputed data, we constructed PGIs for several phenotypes and evaluated the difference in their predictive power. Below, we outline these quality control and imputation procedures.

***Pre-imputation variant and sample quality control.*** We started by filtering out 115,031 variants with minor allele frequency less than 1%, 106,445 variants with genotyping call rate less than 90% and 68,474 variants with Hardy-Weinberg equilibrium exact test *P* value less than $10^{-4}$. Some variants in the data had identifiers of the form "GSA-rs#". We renamed these variants by removing the "GSA-" prefix. This resulted in 64 pairs of duplicate rs-numbers. We removed the first occurring variant from each pair, which left us with 398,604 variants.

Next, we conducted the mild and strict sample-level QC. In the strict-QC pipeline, we dropped 260 individuals whose genotype missingness rate was more than 20% within any chromosome, 14 individuals with sex mismatch identified by comparing reported and inferred sex, and 59 individuals with excess heterozygosity/homozygosity identified as those with Plink1.9[3] --het $F$ coefficient less than -1 or greater than 1 (Figure S5). In the mild-QC pipeline, we applied a per-chromosome genotype missingness rate cutoff of 50%, leading to the exclusion of 36 individuals. The same 14 individuals with sex mismatch were removed, followed by 22 heterozygosity/homozygosity outliers with Plink1.9 --het $F$ coefficient less than -1.5 or greater than 1 (Figure S5). 2,324 and 2,541 individuals remained after strict- and mild-QC, respectively.

We filtered out individuals of non-European ancestries from both data sets prior to imputation. We identified non-European-ancestry individuals using the mild-QC data because the two data sets contained the same set of variants, and the strict-QC sample was a subset of the mild-QC sample. To do so, we filtered out 119,486 variants with a call rate less than 95% and pruned the remaining variants using a 1Mb rolling window (incremented in steps of 5 variants) and a $R^2$ threshold of 0.3, leaving 178,447 approximately independent variants. Next, we merged the data with all samples from the third phase of the 1000 Genomes Project[4,5], restricting to the 170,203 variants available in the pruned Gene-SOEP data with MAF > 1% in the merged sample. We calculated the first four principal components (PCs) of the genetic data using the 1000 Genomes subsample and projected the individuals in the Gene-SOEP sample onto these. Finally, we plotted the first four PCs against each other and identified individuals of European ancestries as those that cluster together with the 1000 Genomes EUR sample. Because the average variant call rate in the Gene-SOEP data was lower than ideal even after the 95% call rate filter, the clustering of Gene-SOEP samples with 1000 Genomes populations was not clearly identifiable. As imposing a stricter call rate filter would lead to the exclusion of too many variants which might also introduce a significant amount of noise into the PCs, we decided to use the current PCs but be lenient in the PC cutoffs for identifying individuals of European ancestries (Figure S6) and repeat the PC analysis after imputation using a larger set of SNPs. After the ancestry filtering, 2,299 and 2,497 individuals remained in the strict- and mild-QC data, respectively.

Prior to imputation, we conducted some reformatting steps and additional quality control checks. First, we used the "HRC-1000G-check-bim.pl"[1] script to check our data against the Haplotype Reference Consortium (HRC) reference panel (r1.1)[7] for strand alignment, position, reference allele assignment and frequency differences. Of the 398,604 variants, 8,415 variants with identifiers and chromosome and base pair positions that could not be matched to the HRC reference panel, 669 indels, 327 palindromic single nucleotide polymorphisms (SNPs) with

---

[1] https://www.well.ox.ac.uk/~wrayner/tools/HRC-1000G-check-bim-v4.3.0.zip, accessed on 12 December 2021.

MAF > 0.4, 403 SNPs whose alleles did not match the HRC panel, and 695 duplicates were removed from both the mild- and strict-QC datasets. In addition, 1,050 SNPs with allele frequency difference with the HRC panel exceeding 0.2 were removed from the mild-QC data, leaving 387,045 SNPs. In the strict-QC data, 1,042 such SNPs were identified, leaving 387,053 SNPs. In both datasets, 30,366 SNPs that were reported on a different strand compared to the HRC reference panel were flipped. Finally, the reference/alternative allele assignment for 388,498 SNPs was switched to match the HRC reference panel. The data were then converted into per-chromosome vcf files using Plink1.9[3], sorted using vcftools[3,8], and checked against the 1000 Genomes Phase 2 reference genome sequence based on NCBI GRCh37 using the "checkVCF.py"[2] script. We observed no important issues by "checkVCF.py".

***Imputation.*** We imputed the 387,045 SNPs from the mild-QC and 387,053 from the strict-QC datasets separately against the HRC reference panel (r1.1) [7] consisting of 64,940 haplotypes of predominantly European ancestry for 32,470 individuals. The genotypes were split into 154 chunks of 20Mb and for each chunk, QC checks were conducted to ensure data validity prior to phasing and imputation. In the mild-QC data, these checks led to the exclusion of 8 chunks for which at least one sample had a call rate less than 50%. No chunks were excluded from the strict-QC data. The squared correlation of allele frequencies between the reference panel and mild-QC data was 0.996, with 1,027 potential mismatches ($\chi^2 > 300$). In the strict-QC data, the squared correlation was 0.997 and 601 potential mismatches were reported. Phasing was done using Eagle2[9] with 5Mb overlap between chunks. Imputation was conducted using Minimac4[10], with a window of 500kb.

Imputation was completed for 2,497 individuals and 23,185,386 SNPs with imputation accuracy ($R^2$) greater than 0.1 in the mild-QC data, and 2,299 individuals and 22,201,548 SNPs with $R^2$> 0.1 in the strict-QC data. Approximately 66% of these SNPs are rare SNPs with MAF<0.01 (15,456,273 in mild-QC, 14,445,952 in strict-QC), ~10% are low-frequency SNPs with 0.01≤MAF<0.05 (2,266,025 in mild-QC, 2,296,334 in strict-QC) , and the remaining ~24% are common SNPs (5,463,110 in mild-QC, 5,463,110 in strict-QC).

The average imputation accuracy in the mild-QC data is 0.664, with 6,501,748 SNPs having $R^2 < 0.5$. In the strict-QC data, the average imputation accuracy is 0.695 with 5,502,428 SNPs below an $R^2$ of 0.5. Figure S7 and S8 show the MAF and $R^2$ distributions of the mild- and strict-QC datasets. Figure S9 shows the mean $R^2$ by MAF bins of width 0.01. As expected, the mean $R^2$is much lower for rare SNPs (0.545 in mild-QC, 0.578 in strict-QC) compared to low-frequency (0.864 in mild-QC, 0.879 in strict-QC) and common SNPs (0.918 in mild-QC, 0.927 in strict-QC).

### 4. Ancestry

In order to make sure that there are no remaining individuals of non-European ancestries in the samples that we were unable to identify with only genotyped SNPs due to the low data quality,

[2] https://github.com/zhanxw/checkVCF, accessed on 12 December 2021

we repeated the ancestry filtering step using imputed data. We merged the mild- and strict-QC datasets separately with all samples from the third phase of the 1000 Genomes Project[7], restricting to HapMap3 SNPs[11] with genotyping call rate greater than 99% and MAF>1% in the merged sample (1,210,703 SNPs in mild-QC, 1,210,959 in strict-QC). The remaining steps were identical to the ancestry filtering with genotyped SNPs described in Section 3. In line with the QC steps before, we chose more lenient PC cutoffs for the mild-QC dataset and filtered out two individuals identified to be of non-European ancestries. From the strict-QC data, we filtered out 37 individuals. Figures S10 and S11 show the PC plots for the mild- and strict-QC data, respectively.

### 5. Genetic principal components

Principal components of the genetic data have been shown to accurately model genetic ancestry differences across individuals and are used as controls against spurious associations between genetic variables and environmental outcomes that may result from population stratification (i.e. systematic differences in allele frequencies of SNPs due to genetic ancestry correlating with differences in environments across ancestries).[12] In genetic analyses, it is standard to control for at least the first 10 PCs, and in analyses where gene-by-environment interactions are being tested, additionally for the interactions between the PCs and the “environment” variable.[13,14] To facilitate analyses that sufficiently account for population stratification, we make the first 20 PCs of the genetic data available to users.

We followed the pipeline outlined in Becker et al.[14] to construct genetic PCs. Briefly, we restricted the mild- and strict-QC samples to individuals of European genetic ancestries as described above and removed SNPs with imputation accuracy < 70% or MAF < 1%, as well as SNPs in long-range LD blocks (chr5:44mb-51.5mb, chr6:25mb-33.5mb, chr8:8mb-12mb, chr11:45mb-57mb). We pruned the remaining SNPs using an $r^2$ cutoff of 0.01 and a 1Mb rolling window incremented in steps of 5 variants in Plink1.9[3], resulting in 162,306 and 159,312 approximately independent SNPs in mild- and strict-QC data, respectively. Using these sets of SNPs, we estimated a genetic relatedness matrix for each dataset, calculated the first 20 PCs in the sample of unrelated individuals (Plink1.9 genetic relatedness coefficient < 0.05; 1,983 in mild-QC data, 1,905 individuals in strict-QC), and projected the remaining individuals onto these PCs.

### 6. Genetic family relationships

We used KING[15] to infer biological family relationships among the participants. As it was done for the genetic ancestry analysis, we relied on the imputed SNP data and used mild-QC data to include as many individuals as possible. Following the recommendation by the authors of the software, we applied a minimal level of SNP filters and used 1,159,581 HapMap3[11] SNPs with imputation accuracy > 70%, genotyping call rate > 99%, and MAF > 0.1%.

As a result, we identified 5,585 pairs of individuals who are related up to fourth degree (2 identical twins, 510 parent-offspring, 132 full-siblings, 140 second-degree, 75 third-degree,

4,726 fourth-degree). However, only 603 pairs remained when the sample was restricted to individuals whose genotype call rate, computed only from directly genotyped SNPs, was greater than 90% ($N$ = 2,070; 415 parent-offspring, 103 full-sibling, 56 second-degree, 20 third-degree, 9 fourth-degree). This difference suggests that most of the genetically inferred distant relatives are likely to be incorrect due to low data quality.

When considering only the individuals whose genotyping call rate was greater than 90% using directly genotyped SNPs, 97% of the pairs in the SOEP-IS have consistent self-reported and genetic family relationships. We found that most of the remaining inconsistencies are due to self-reported full siblings who are likely to be only half siblings (13 out of 97 pairs). We also found eight self-reported parent-child pairs that are non-biological.

Furthermore, restricting to the individuals with the genotype call rate greater than 90%, we identified 88 pairs whose family relationship information was not available in the survey data. These pairs consist of 7 parent-offspring, 19 full-sibling, 33 second-degree relative, and 29 third- or fourth-degree relative pairs.

Overall, out of 2,497 individuals, we genetically identified 703 individuals with at least one first-degree relatives (parent-child or full sibling) and 728 individuals that have at least one relative with at least third-degree of relatedness (first cousins or great grandparent-child). 1,769 individuals do not have close relatives on the basis of the genetic data. Note that the related pairs reported here are not mutually exclusive and some individuals can be related to multiple people.

Figure S12 summarizes the quality control process we applied to the genetic data.

### 7. Polygenic indices

A “polygenic index” (PGI, also known as polygenic score) is a DNA-based predictor capturing the relationship between a large number of SNPs from an individual’s genome and a phenotype such as disease risk or a social or behavioral outcome. Oftentimes, PGIs are constructed to predict complex traits that are influenced by many genetic markers with small effects, such as height or educational attainment. For example, Lee et al. [16] use ~1.2 million HapMap3[11] SNPs to construct a PGI for educational attainment (EA) which explains ~11% of the variance in EA in independent samples. A height PGI based on Yengo et al.[17] constructed using the same set of SNPs explains up to 32% of the variance in height.[14]

A PGI is obtained as a weighted sum of a person’s genotypes at a number of genetic markers. Methodologies for PGI construction differ mainly across two dimensions: (i) the weights, and (ii) the set of genetic markers included in the PGI. A common strategy is to set the weights equal to the coefficient estimates from a genome-wide association study (GWAS) conducted in a sample independent from the prediction sample. GWAS are hypothesis-free studies that test the association between a phenotype and each of millions of genetic markers in separate univariate regressions. Since genetic markers in proximity are correlated with each other (i.e. “linkage disequilibrium” or LD), including all available genetic markers in a PGI would result in double-counting genetic variants that are in high LD with each other, which decreases the predictive accuracy of the resulting PGI. Therefore, pruning the GWAS summary statistics to obtain a set of approximately independent markers to include in the PGI is common practice. Markers are usually also filtered for the association $P$ value that they have with the phenotype so that only the markers that are most likely to be truly associated with the phenotype are included in the PGI. Although this methodology, called "Clumping and Thresholding (C+T)"[18,19] is often used due to its simplicity, it has certain disadvantages. If the pruning (or clumping) is done too strictly, relevant genetic markers may get removed. If it is done too liberally, the double-counting problem persists. As a solution, new methodologies have been developed that use Bayesian approaches to adjust the GWAS coefficient estimates for LD, nullifying the necessity to include independent markers in the PGI. PGIs made using these methodologies have been shown to be more predictive than their C+T counterparts.[20–23]

The PGI Repository[14] by the Social Science Genetic Association Consortium (SSGAC) uses one such Bayesian methodology, LDpred[20], to construct PGIs for 47 phenotypes in several datasets. Their PGIs are made using the largest GWAS available at the time of publication for almost all of the 47 phenotypes, obtained by meta-analyzing summary statistics from multiple sources, including several novel large-scale GWASs conducted in UK Biobank[24] and the personal genomics company 23andMe. We used the same pipeline and GWAS summary statistics to construct the 47 PGIs from the Repository in the SOEP-IS data, as well as for 19 additional medical outcomes for which well-powered GWAS summary statistics were available (see Table 3 in the main text). In contrast to the pipeline employed by Becker et al., we did not use the E($R^2$)>1% threshold for including these medical outcomes.

Prior to making the PGIs for all 66 phenotypes, we conducted analyses to compare the predictive power of PGIs made using mild- and strict-QC data for height and BMI. For both phenotypes, we estimated three quantities: (i) the increase in the coefficient of determination ($R^2$) when the strict-QC PGI for the phenotype is added to a regression of the phenotype on controls (year of birth, year of birth$^2$, sex, interactions between the sex and age terms, genotype batch indicators, and the top 20 genetic PCs), i.e. the "incremental-$R^2$" of the strict-QC PGI (ii) the incremental-$R^2$of the mild-QC PGI, and (iii) the incremental-$R^2$ of the mild-QC PGI when the estimation sample is restricted to the individuals available in the strict-QC sample. Since height and BMI were surveyed multiple times across the waves, we first residualized height and BMI for age, age$^2$, sex and their interactions within each wave and took the mean for each individual; then, as covariates, we used only genotype batch indicators and the top 20 genetic PCs. We obtained 95% confidence intervals by bootstrapping the sample 2,000 times (Figure 3 in the main text). For this analysis, we only included unrelated individuals by removing one person from each related pair up to the third degree of relatedness. The related pairs here were restricted to those with matching self-reported and genetically inferred relationships and those genetically identified where both individuals had genotyping call rate greater than 0.9.

In general, the strict-QC PGIs performed slightly better than the mild-QC PGIs, but the differences were not statistically distinguishable (Supplementary Table 2). Also, the predictive power of the mild-QC PGIs is very close to that of the strict-QC PGIs when the samples are restricted to the individuals in the strict-QC data. These results suggest that the inclusion of low-quality samples in the mild-QC imputation did not have a big impact on the quality of imputation, at least for HapMap3 SNPs[11], which the PGIs are based on. Therefore, we decided to use the mild-QC data to construct the PGIs for all 66 phenotypes.

The data that we will publicly share includes 20 genetic PCs, all constructed PGIs, as well as indicators for individuals that did not pass the strict-QC pipeline. This allows users to decide whether they prefer to conduct their analyses using the full sample for which PGIs were constructed or the slightly smaller set that passed strict QC. The choice between these two options may depend on statistical power considerations and the specific research question being asked.

Below, we summarize the main steps in the PGI Repository pipeline and explain how they were applied to the SOEP-IS data.

In the PGI Repository, there are two types of PGIs: Single-trait and multi-trait. Single-trait PGIs are regular PGIs based on univariate GWAS analyses. Multi-trait-PGIs are based on multivariate analyses of a phenotype and its supplementary phenotypes, defined as those phenotypes (if any) whose genetic correlation with the target phenotype exceeds 0.6 in absolute value, conducted using the MTAG software.[25] All single- and multi-trait PGIs for the 47 phenotypes from the Repository have expected out-of-sample predictive power greater than 1% as this was the criterion for inclusion that Becker et al.[14] implemented. For the remaining 19 phenotypes, we only constructed single-trait PGIs and did not implement an inclusion criterion based on expected predictive power.

All PGIs were constructed in Plink2[3,26] using imputed genotype probabilities for HapMap3 SNPs[11]. We adjusted the SNP weights for LD using LDpred[20], assuming a fraction of causal SNPs equal to 1. LDpred requires a reference genotype dataset to estimate the correlation structure between SNPs. As in Becker et al., we used the public release of the HRC Reference Panel (r1.1)[7] after applying several quality control filters to estimate LD patterns, with a LD window equal to the number of SNPs common between LD reference data, Gene-SOEP mild-QC genotype data, and summary statistics left after LDpred quality control filters, divided by 3,000. Table 3 in the main text lists the 55 phenotypes for which we made single trait PGIs, the number of SNPs included in each PGI, and the GWAS sample sizes of the summary statistics that the PGIs are based on, the source of the summary statistics. SI Table 3 lists all multi-trait PGIs, the supplementary phenotypes that were used to construct them, as well as the GWAS equivalent sample size reported by MTAG[25] , defined as the GWAS sample size required to obtain a mean $\chi^2$-statistic equal to that attained by the MTAG analysis, which, under the LD score regression assumptions (Bulik-Sullivan et al. 2015), is equal to

$$N_{GWAS-equiv,j} = N_{GWAS,j}\ \frac{\bar{\chi}^2{}_{MTAG} - 1}{\bar{\chi}^2{}_{GWAS} - 1},$$

where $N_{GWAS,j}$ is the sample size of SNP $j$ in the univariate GWAS results for the phenotype, $\bar{\chi}^2{}_{GWAS}$ is the mean $\chi^2$ $-$statistic in the univariate GWAS results, and $\bar{\chi}^2{}_{MTAG}$ is the mean $\chi^2$ $-$statistic in the MTAG results.
More information on how the PGIs were constructed can be found in Becker et al[14].

### 8. Author contributions

Designed and oversaw the study: Philipp Koellinger, Ralph Hertwig, Gert Wagner
Data collection: Philipp Koellinger, Gert Wagner, Bettina Maria Zweck, Jan Goebel, David Richter
Data quality control and imputation: Aysu Okbay, Hyeokmoon Kweon
Constructed genetic principal components: Aysu Okbay
Constructed polygenic indices: Aysu Okbay, Annemarie Schweinert
Contributed computer scripts and feedback: Richard Karlsson Linnér
Data analysis: Annemarie Schweinert, Hyeokmoon Kweon, Lisa Reiber, David Richter, Bettina Maria Zweck
Figures and tables: Aysu Okbay, Annemarie Schweinert, Hyeokmoon Kweon, David Richter, Jan Goebel
Wrote manuscript: Philipp Koellinger, Annemarie Schweinert, Hyeokmoon Kweon, Aysu Okbay
Wrote supplementary information: Aysu Okbay, Hyeokmoon Kweon, Philipp Koellinger, Annemarie Schweinert
Prepared data for public sharing: David Richter, Aysu Okbay, Hyeokmoon Kweon, Annemarie Schweinert, Bettina Maria Zweck, Lisa Reiber
Project administration: Philipp Koellinger
Funding acquisition and resources: Ralph Hertwig, Philipp Koellinger, K. Paige Harden, Gert Wagner, Daniel Belsky, Pietro Biroli, Rui Mata, Elliot Tucker-Drob
Edited manuscript and supplementary information: All co-authors

### 10. Supplementary Figures

**Figure S1.** Histogram of minor allele frequencies of genotyped autosomal biallelic SNPs

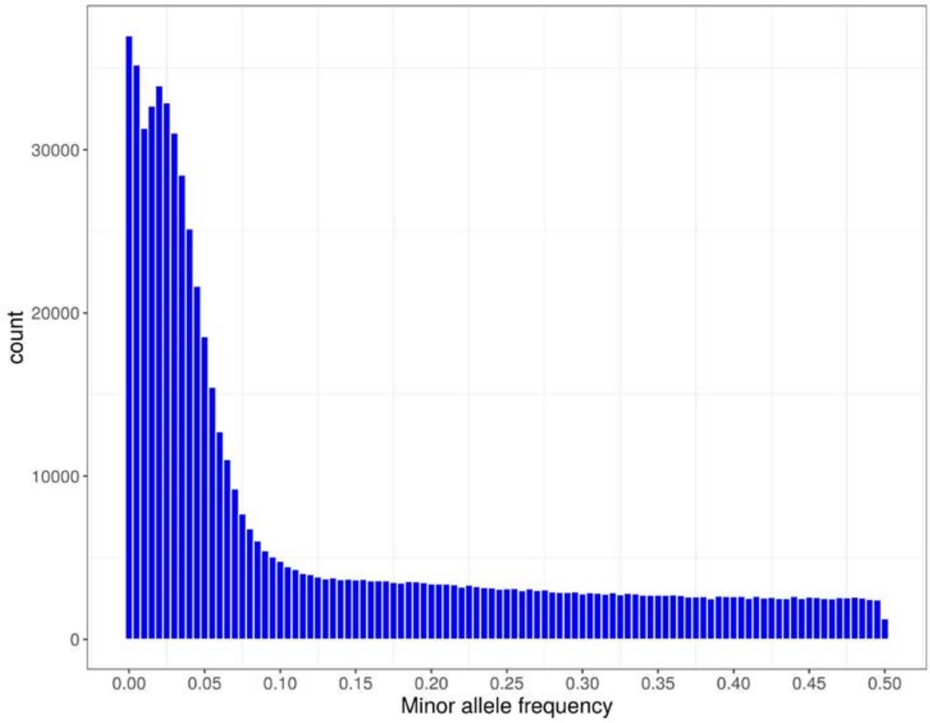

**Figure S2.** Histogram of genotype call rates of SNPs and samples

**A.** SNP call rates

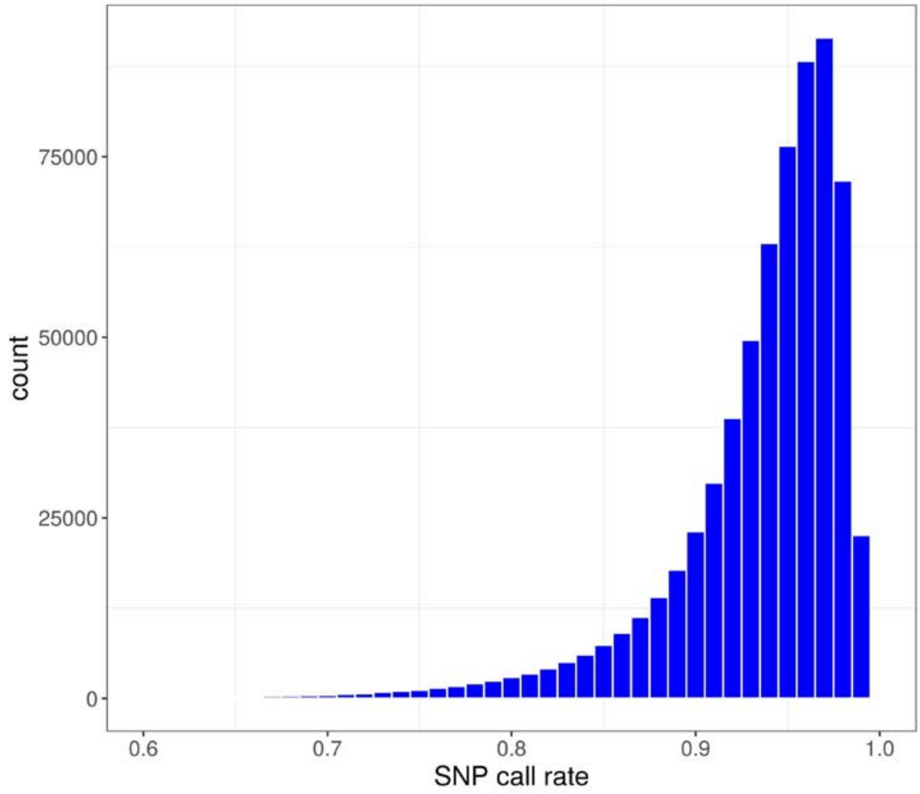


**B.** Sample call rates

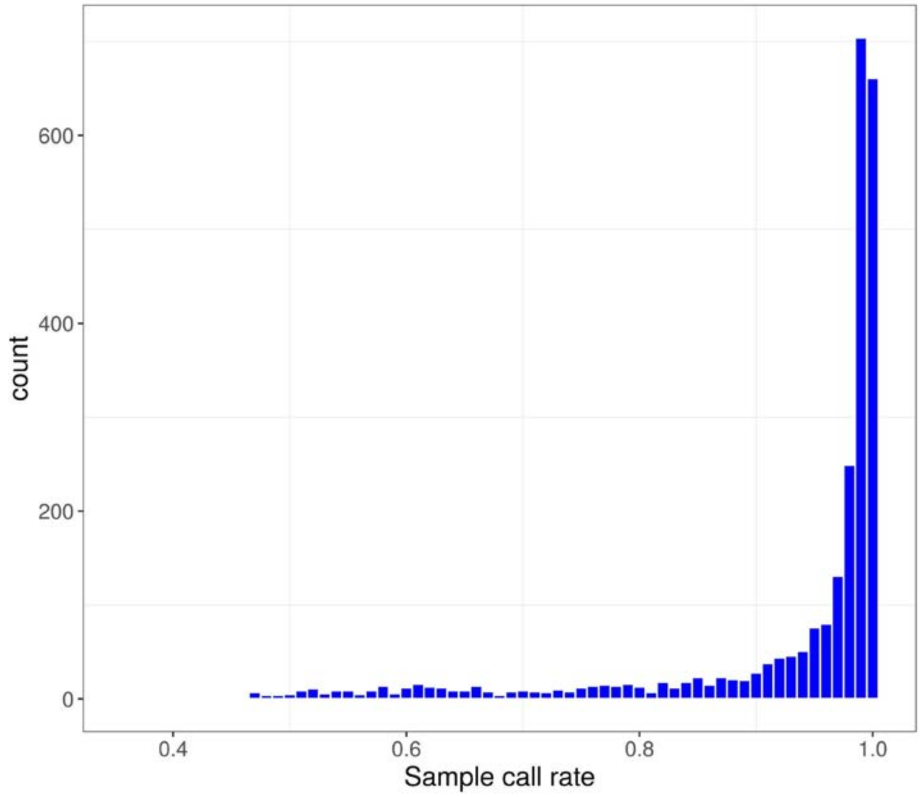


Note: only autosomal biallelic SNPs were used.

**Figure S3.** Binned scatter plot of SNP missing rates over minor allele frequencies

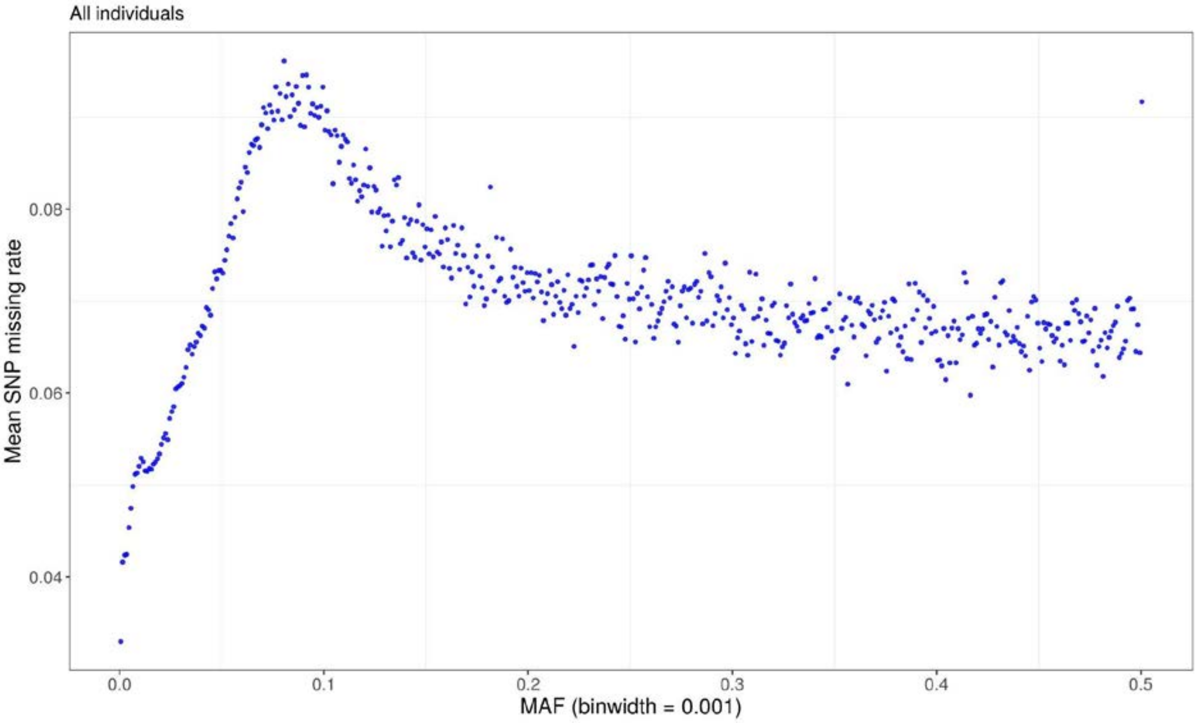


**Figure S4.** Histogram of mean sample call rates by interviewer

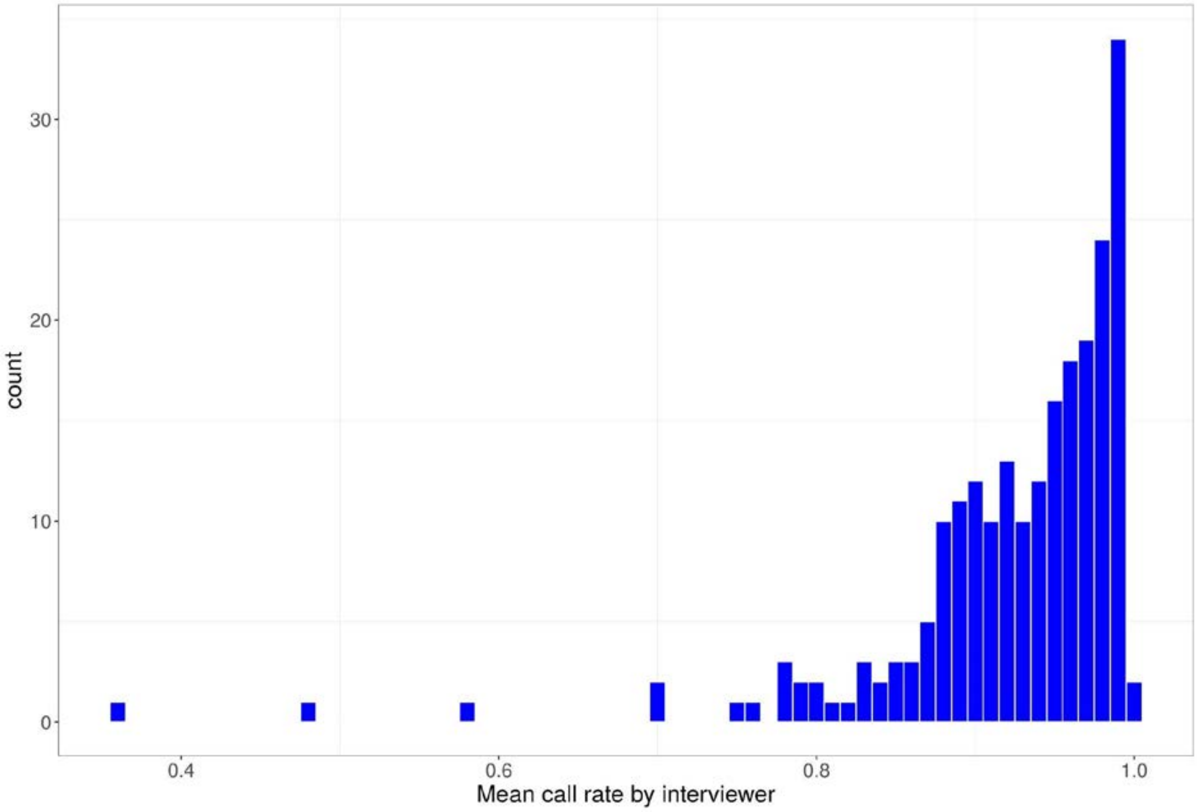

**Figure S5.** Homozygosity / Heterozygosity outliers

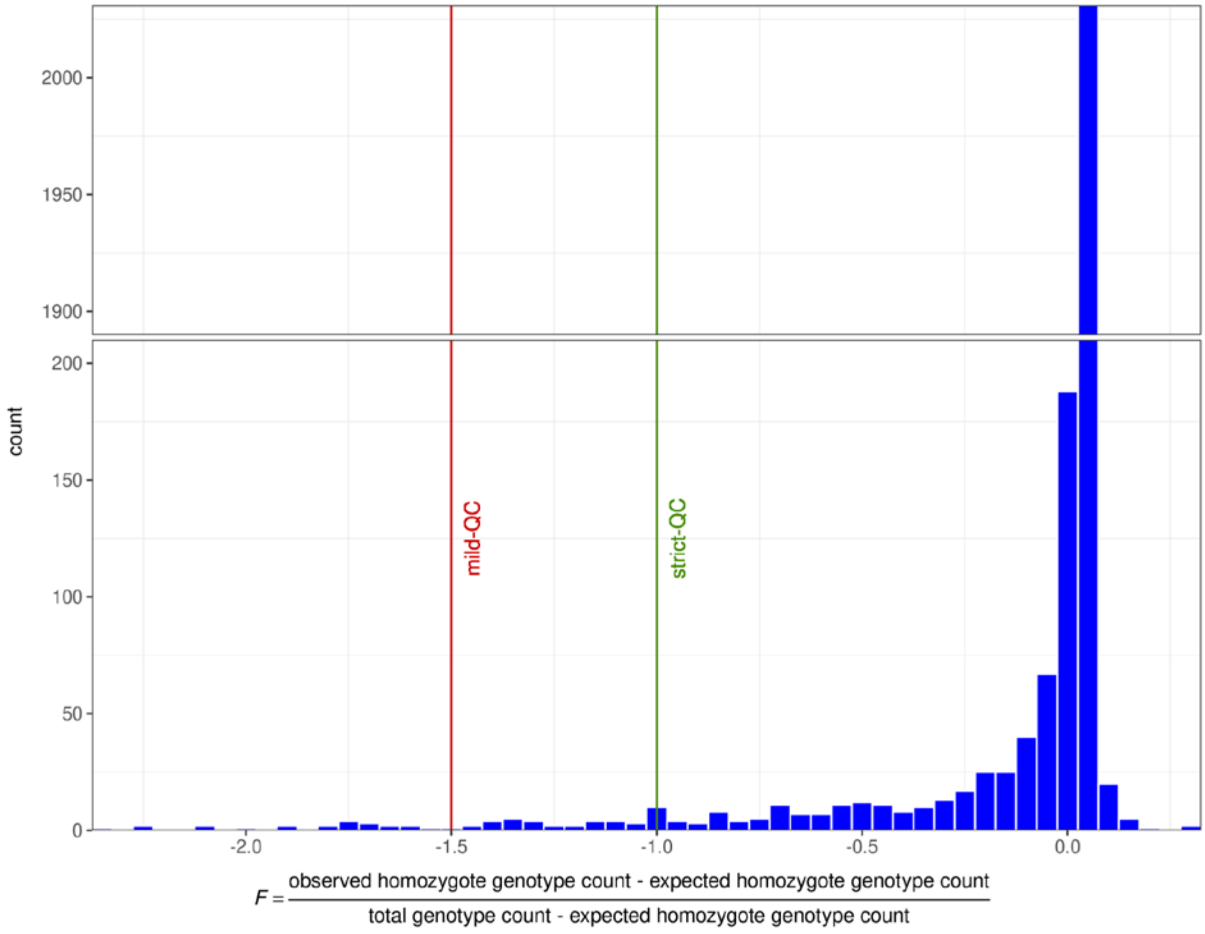

**Figure S6.** Pre-imputation (mild-)QC - Ancestry filtering

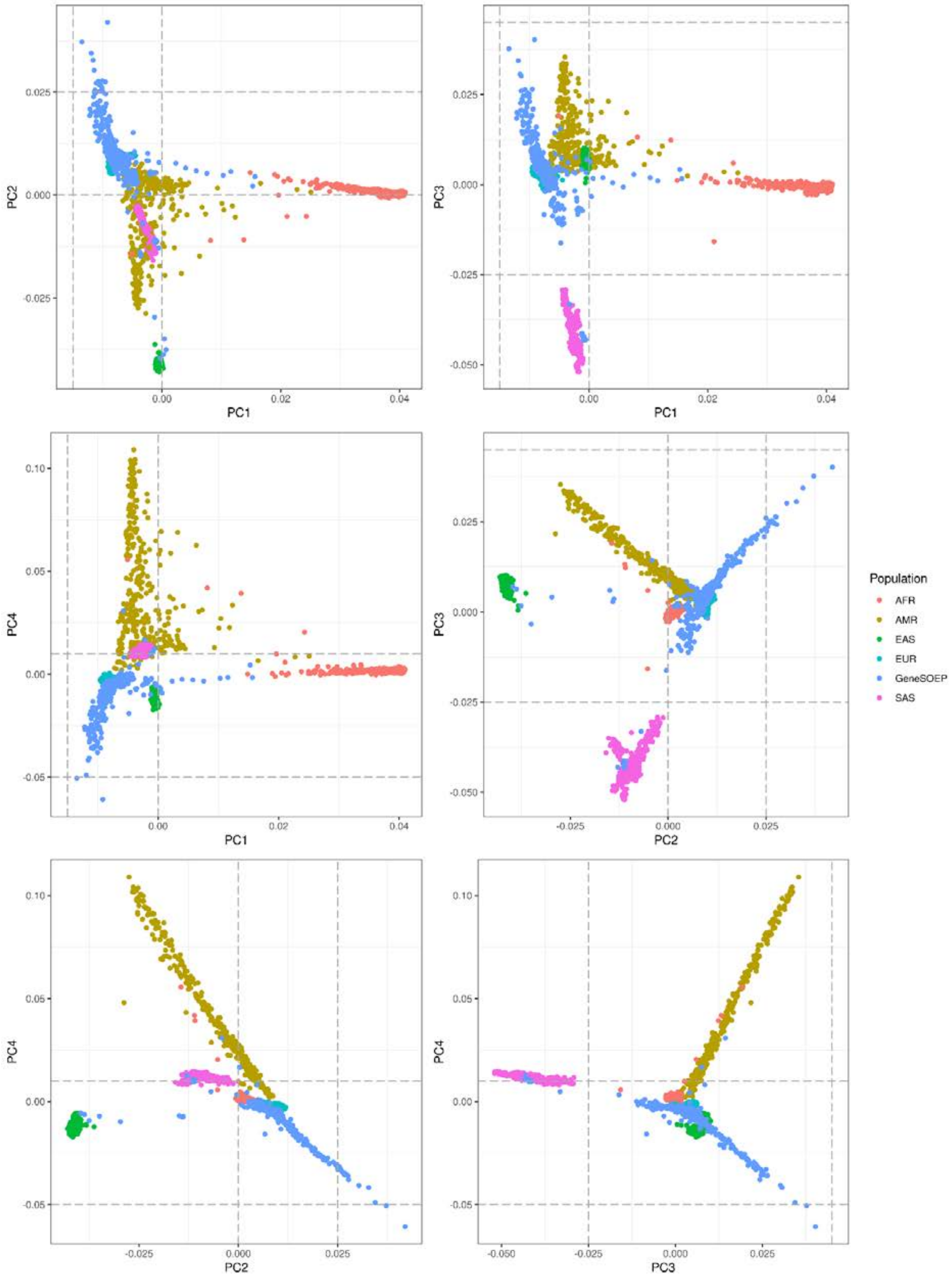

**Figure S7.** Imputed data MAF distribution

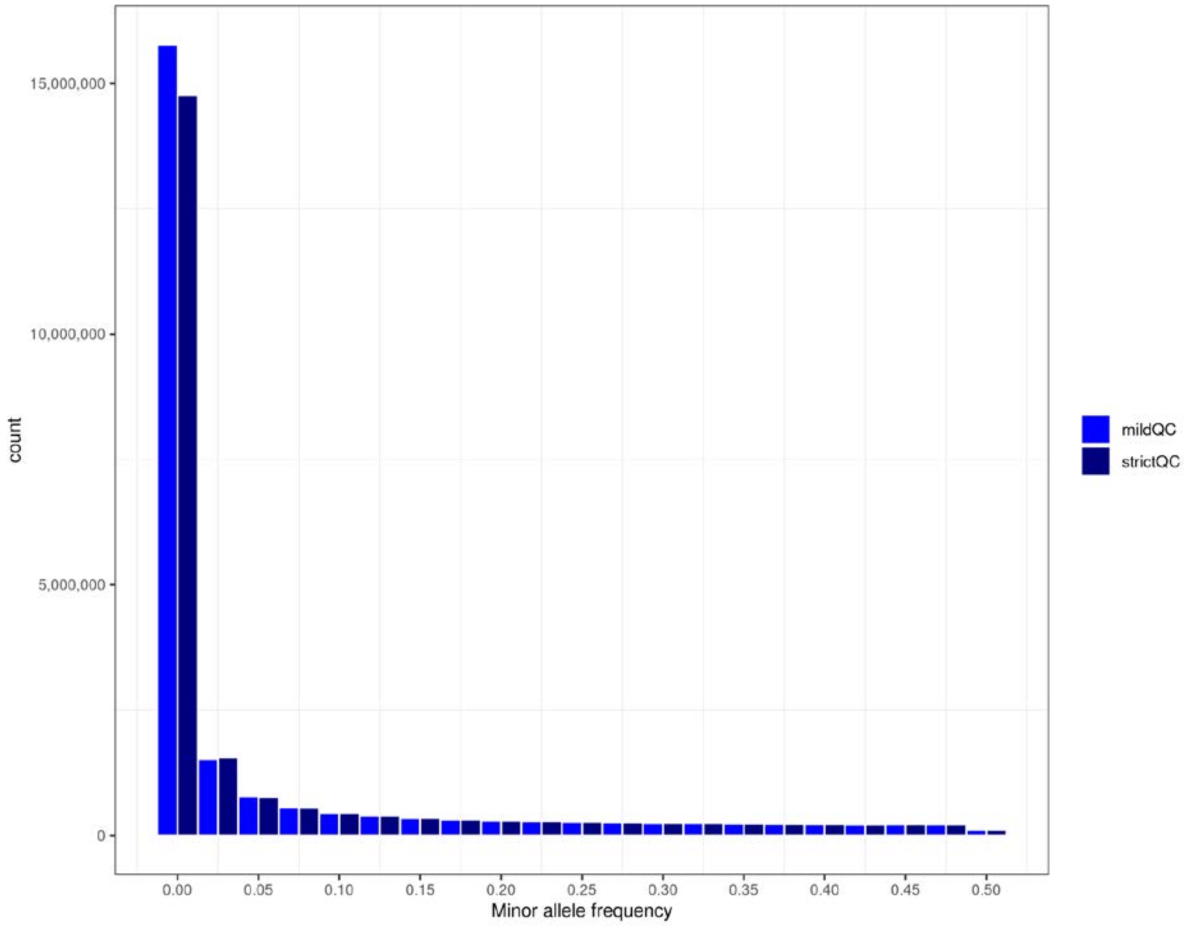

**Figure S8.** Imputation accuracy distribution

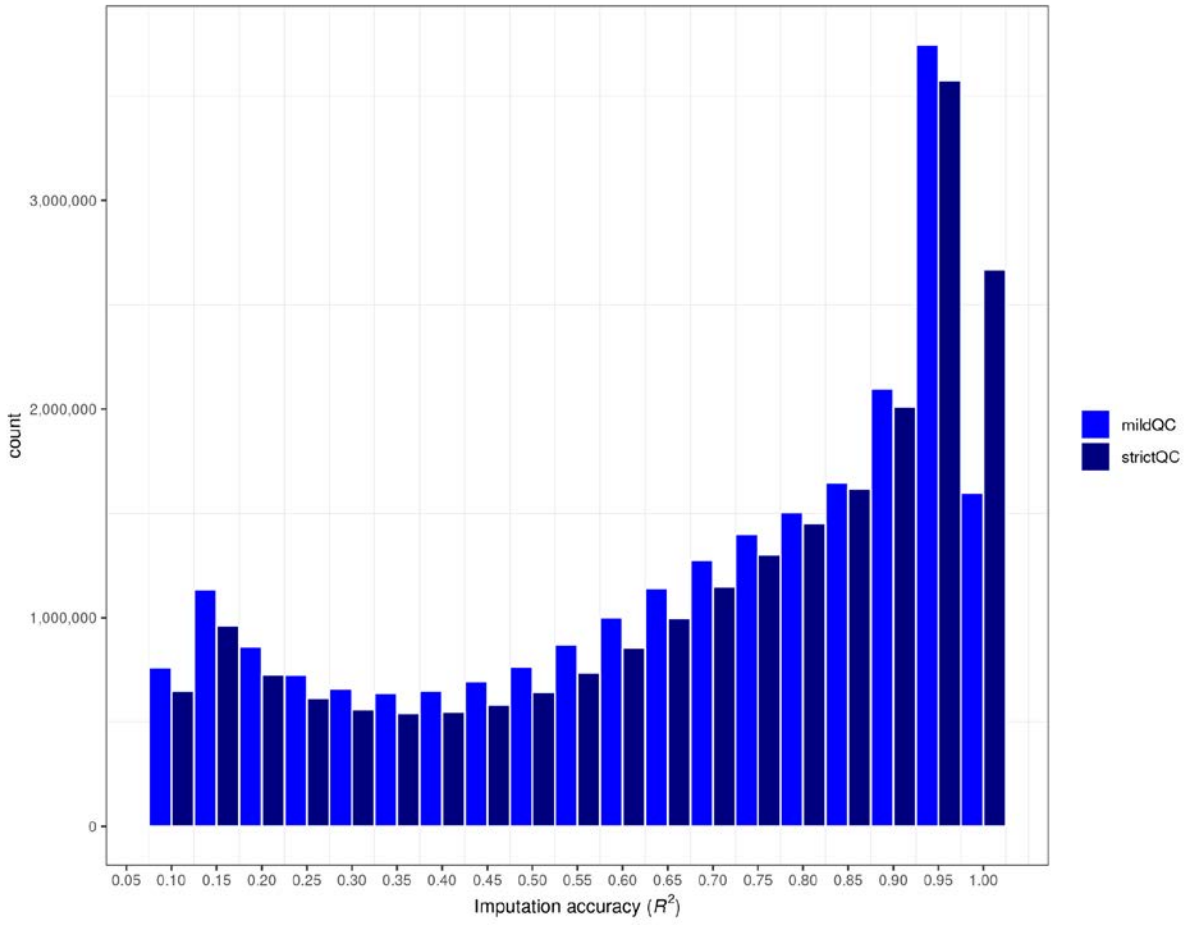

**Figure S9.** Mean imputation accuracy by MAF

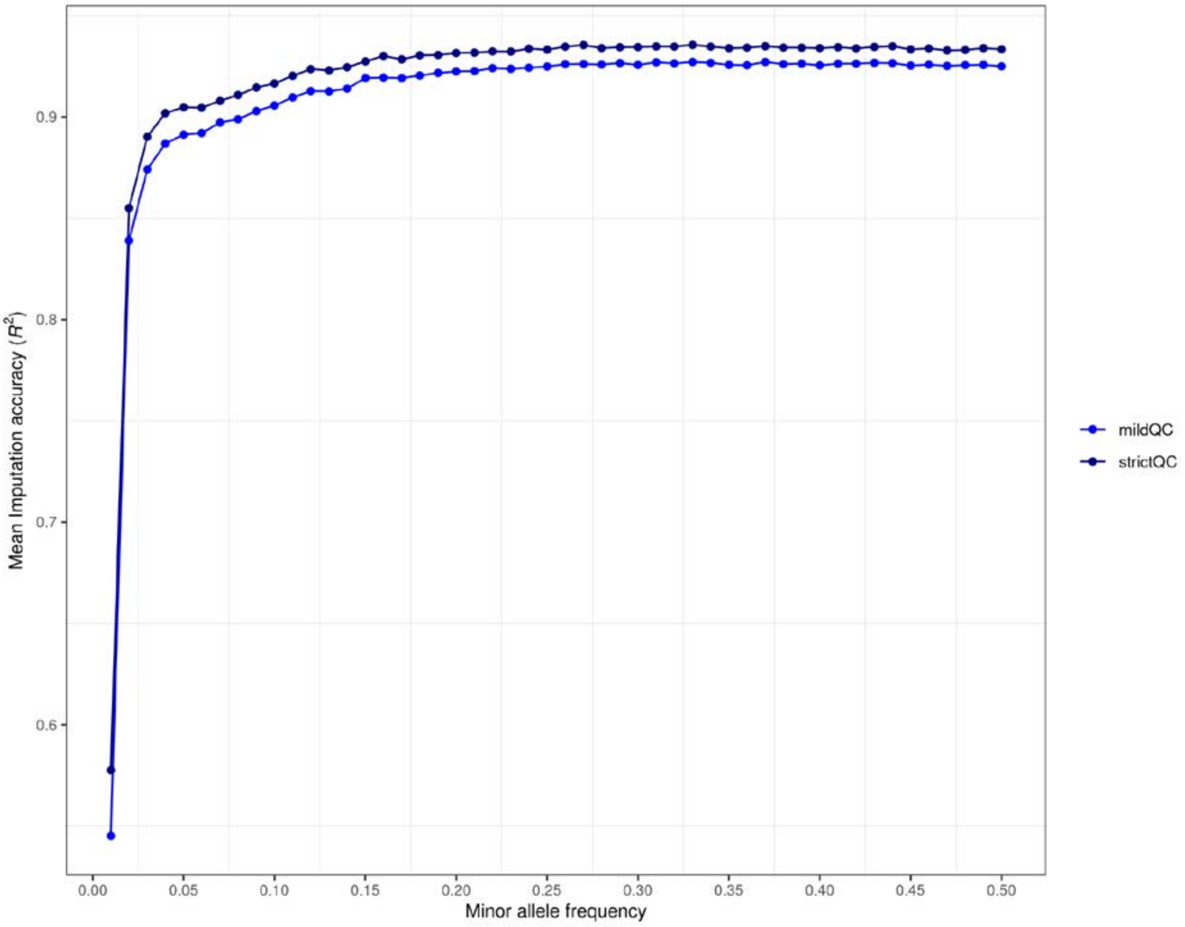

**Figure S10.** Post-imputation mild-QC - Ancestry filtering

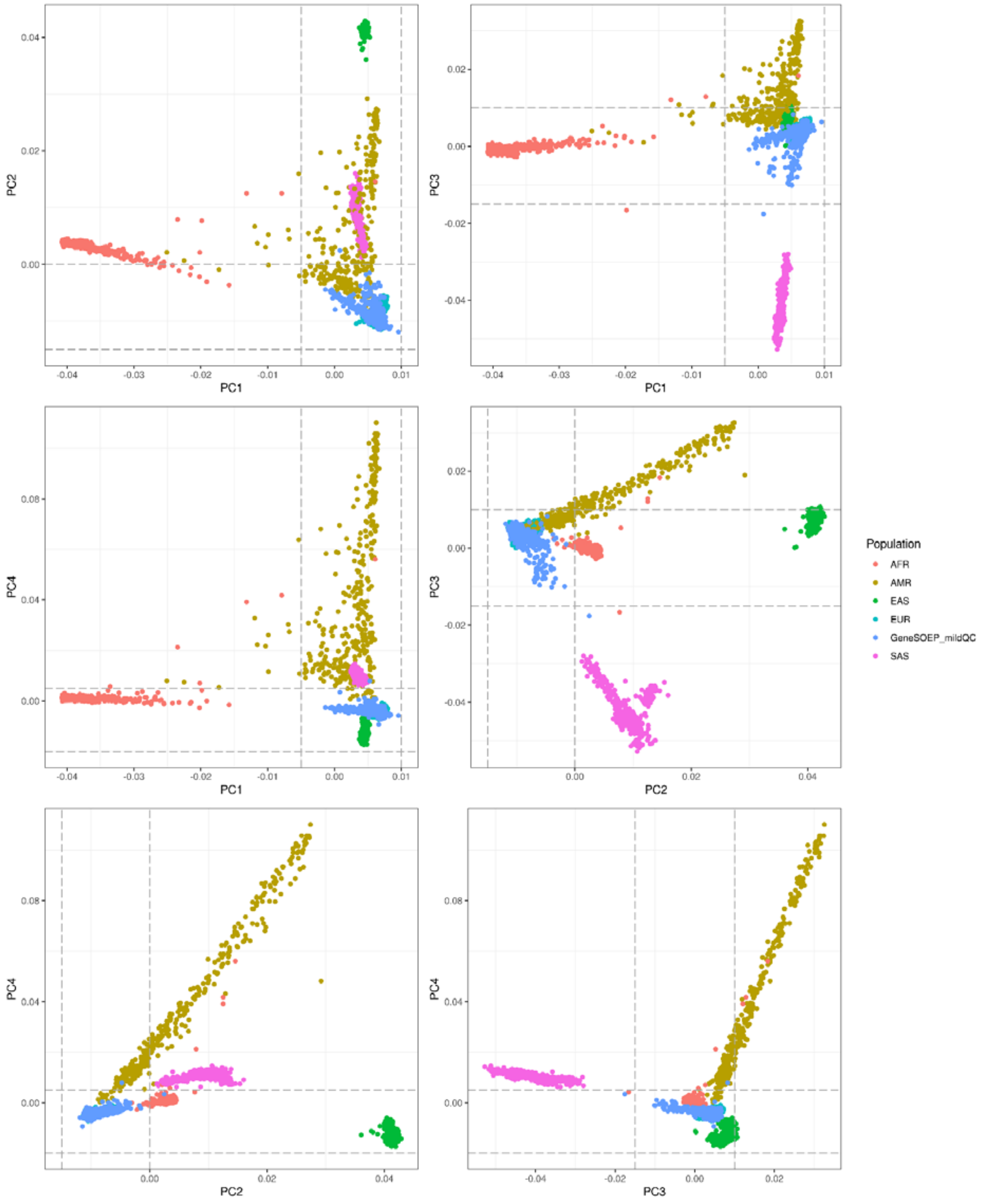

**Figure S11.** Post-imputation strict-QC - Ancestry filtering

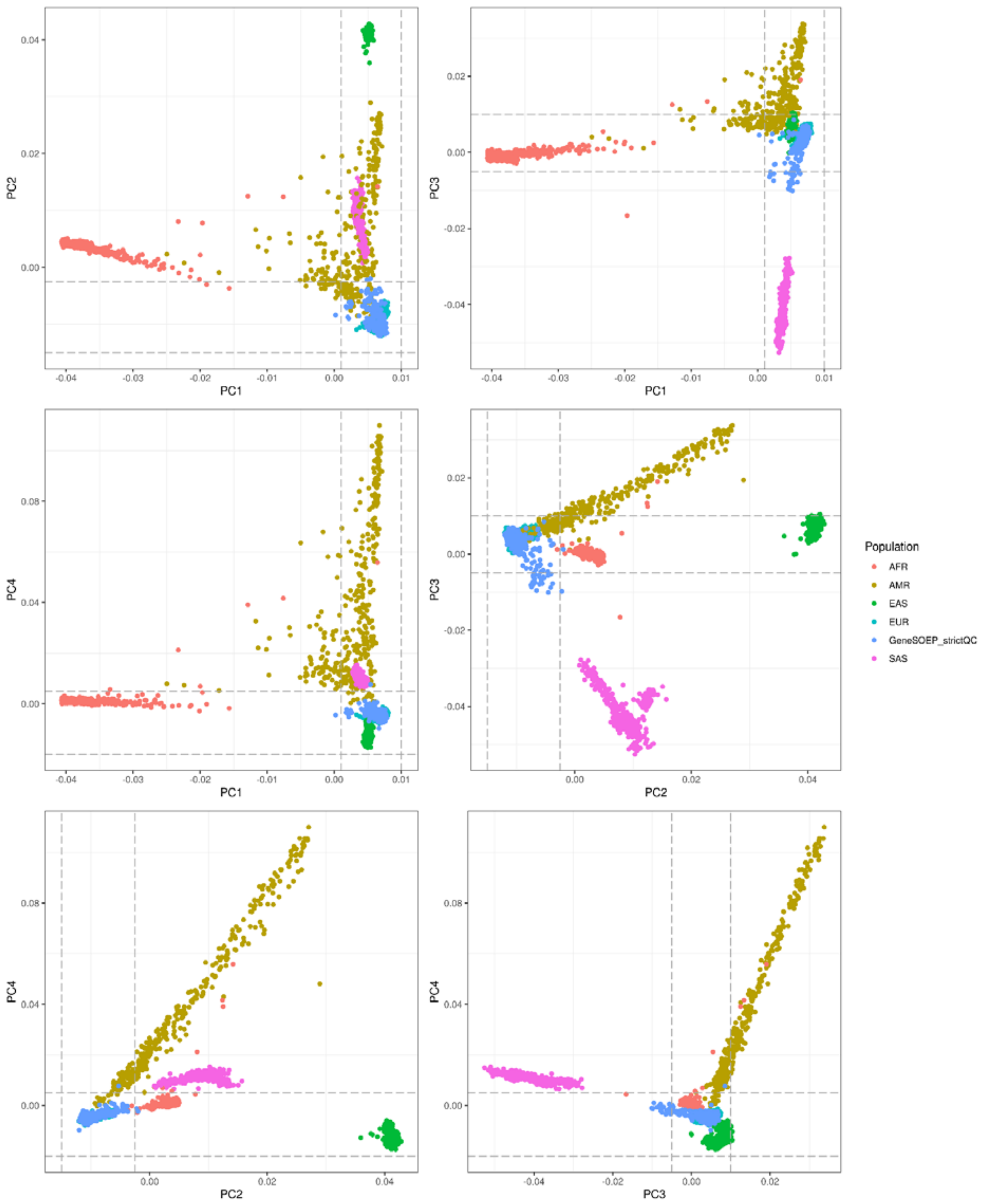

**Figure S12.** Flow-chart quality control of genetic data

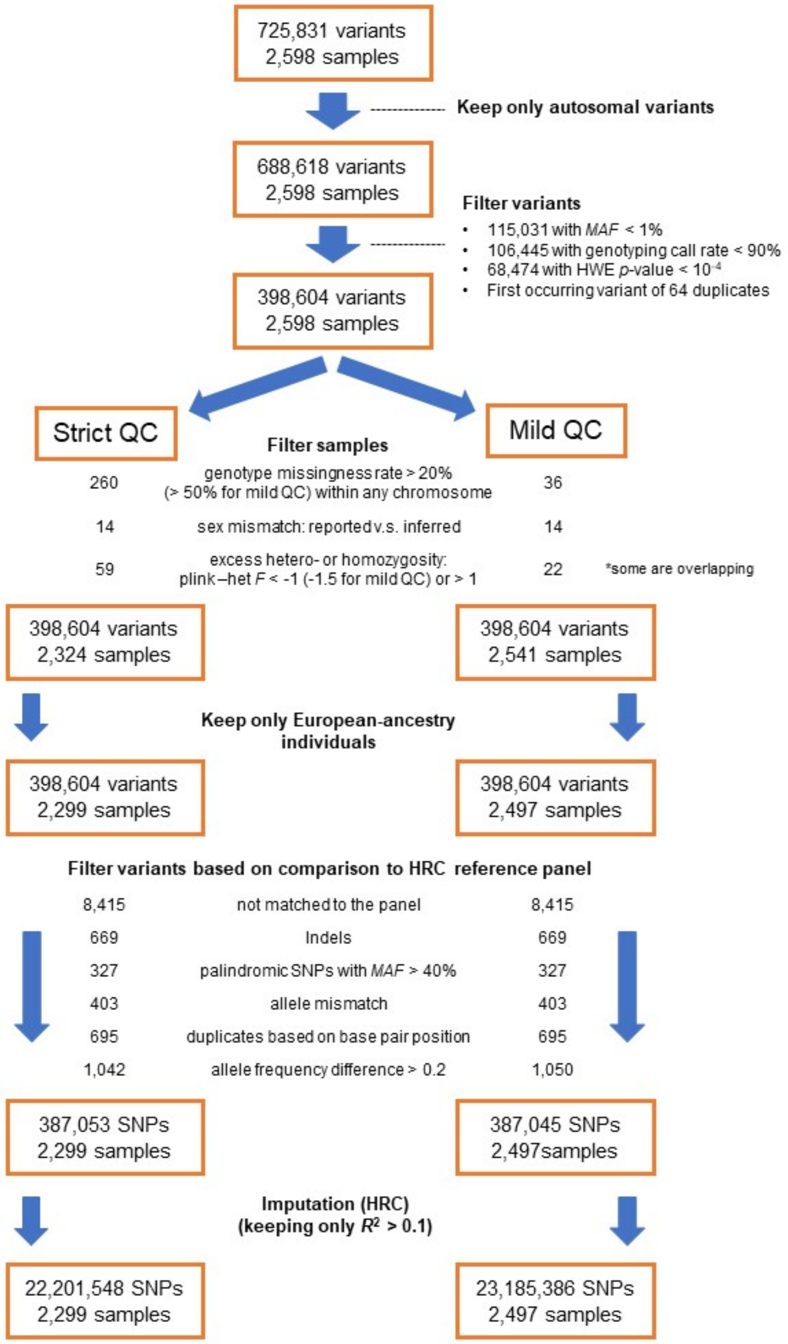
