## Supplementary Tables for "Cohort Profile: Genetic data in the German Socio-Economic Panel Innovation Sample (Gene-SOEP)"

**Index of Supplementary Tables 1 (ordered by appearance in the main text)**

| Table | Contents |
| --- | --- |
| Supplementary Table 1 | Comparison of reported and genetically inferred family relationships |
| Supplementary Table 2 | Polygenic prediction accuracy for height, BMI, and educational attainment |
| Supplementary Table 3 | MTAG polygenic indices |
| Supplementary Table 4 | Predictive accuracy of polygenic indices on self-rated health |
| Supplementary Table 5 | Observed height birth cohort effect estimates |
| Supplementary Table 6 | Observed BMI birth cohort effect estimates |

**Supplementary Table 1: Comparison of reported and genetically inferred family relationships**

Panel A. No sample filter

| | | Genetically inferred relationship | | | | | | |
|---|---|---|---|---|---|---|---|---|
| | | **2nd degree** | **3rd degree** | **4th degree** | **Full sibling** | **Parent-offspring** | **Unrelated** | **Sum** |
| **Reported or inferred relationship from survey** | **2nd degree** | 12 | 1 | 0 | 0 | 0 | 4 | 17 |
| | **Full sibling** | 27 | 6 | 2 | 107 | 0 | 0 | 142 |
| | **Parent-offspring** | 33 | 10 | 13 | 0 | 501 | 52 | 609 |

Panel B. Individuals whose genotype call rate > 90%

| | | Genetically inferred relationship | | | |
|---|---|---|---|---|---|
| | | **2nd degree** | **Full sibling** | **Parent-offspring** | **Unrelated** |
| **Reported or inferred relationship from survey** | **2nd degree** | 8 | 0 | 0 | 4 |
| | **Full sibling** | 13 | 84 | 0 | 0 |
| | **Parent-offspring** | 0 | 0 | 409 | 28 |

**Notes**: Panel A compares the reported family relationships to genetically inferred family relationships. Panel B reports the results only from individuals whose genotype call rates exceed 90% (computed from directly genotyped SNPs). The genetic relationships were estimated with King

**Supplementary Table 2: Polygenic prediction accuracy for height, BMI, and educational attainment**

| | Mild QC | | | Strict QC | | | Mild QC samples in strict QC | | |
|---|---|---|---|---|---|---|---|---|---|
| **Phenotype** | **Estimate (*SE*)** | **$R^2$ (%)** | ***N*** | **Estimate (*SE*)** | **$R^2$ (%)** | ***N*** | **Estimate (*SE*)** | **$R^2$ (%)** | ***N*** |
| Height | 3.871 (0.154) | 22.00 [20.22, 26.41] | 2,094 | 3.791 (0.149) | 23.92 [22.38, 28.67] | 1,904 | 3.782 (0.150) | 23.76 [22.22, 28.50] | 1,904 |
| BMI | 1.861 (0.112) | 11.51 [9.47, 14.76] | 2,086 | 1.859 (0.110) | 12.67 [10.86, 15.96] | 1,897 | 1.876 (0.111) | 12.78 [11.07, 16.07] | 1,897 |
| Educatioanl attainment | 0.853 (0.058) | 8.94 [7.20, 11.71] | 2,036 | 0.857 (0.061) | 9.02 [7.24, 11.81] | 1,857 | 0.880 (0.061) | 9.40 [7.56, 12.27] | 1,857 |

**Notes**: The results are reported separately for mild and strict QC samples as well as the mild QC samples that passed strict QC. Coefficient estimates are measured in standard deviation units of the polygenic index, with standard errors reported in parentheses. Boostrapped 95% confidence intervals for the predictive accuracy measured in $R^2$ are reported in the brackets.

**Supplementary Table 3: Multi-trait polygenic indices in the Gene-SOEP sample**

| Phenotype | # SNPs | GWAS-equivalent *N* | Supplementary Phenotypes | References |
|---|---|---|---|---|
| Adventurousness | 1,070,162 | 800,552 | Risk Tolerance | 1, 17 |
| Age First Birth | 943,209 | 373,210 | Number Ever Born (Women), Number Ever Born (Men), Highest Math, Educational Attainment, COPD, ADHD | 1, 5, 6, 10 |
| Age First Menses (Women) | 1,142,017 | 319,522 | Age Voice Deepened (Men) | 1, 7 |
| Age Voice Deepened (Men) | 1,142,019 | 294,997 | Age First Menses (Women) | 1, 7 |
| Alcohol Misuse | 1,137,769 | 444,496 | Drinks per Week | 1, 8, 13 |
| Allergy - Cat | 1,102,361 | 366,535 | Hayfever, Asthma, Asthma/Eczema/Rhinitis, Allergy - Pollen, Allergy - Dust | 1, 9, 25 |
| Allergy - Dust | 1,102,361 | 356,879 | Allergy - Cat, Hayfever, Asthma, Asthma/Eczema/Rhinitis, Allergy - Pollen | 1, 9, 25 |
| Allergy - Pollen | 1,102,361 | 254,386 | Allergy - Cat, Hayfever, Asthma, Asthma/Eczema/Rhinitis, Allergy - Dust | 1, 9, 25 |
| Asthma | 1,102,426 | 559,863 | Allergy - Cat, Hayfever, Asthma/Eczema/Rhinitis, Allergy - Pollen, Allergy - Dust | 1, 9, 25 |
| Asthma/Eczema/Rhinitis | 1,102,415 | 920,692 | Allergy - Cat, Hayfever, Eczema, Asthma, Allergy - Pollen, Allergy - Dust | 1, 9, 25 |
| Attention Deficit Hyperactivity Disorder (ADHD) | 946,589 | 760,838 | COPD, Age First Birth | 1, 6, 10 |
| Cognitive Empathy | 1,149,525 | 55,982 | Agreeableness | 1, 18, 26 |
| Cognitive Performance | 1,135,783 | 343,411 | Self-Rated Math Ability, Highest Math, Educational Attainment | 1, 4, 5 |
| COPD | 943,332 | 1,869,820 | Self-Rated Health, Age First Birth, ADHD | 1, 6, 10 |
| Delay Discounting | 1,143,145 | 445,313 | Highest Math, Educational Attainment | 1, 5, 27 |
| Depressive Symptoms | 867,549 | 1,306,090 | Subjective Well-Being, Neuroticism, Loneliness, Self-Rated Health | 1, 14, 18, 21, 23 |
| Drinks per Week | 1,137,778 | 1,245,225 | Alcohol Misuse | 1, 8, 13 |
| Educational Attainment | 985,887 | 1,295,788 | Religious Attendance, Highest Math, Delay Discounting, Cognitive Performance, Age First Birth | 1, 4, 5, 6, 24 |
| Extraversion | 1,107,023 | 111,464 | Left Out of Social Activity | 1, 18, 19 |
| Hayfever | 1,102,417 | 518,480 | Asthma, Asthma/Eczema/Rhinitis, Allergy - Pollen, Allergy - Dust, Allergy - Cat, Eczema | 1, 9, 25 |
| Highest Math | 985,893 | 801,291 | Self-Rated Math Ability, Educational Attainment, Delay Discounting, Cognitive Performance, Age First Birth | 1, 4, 5, 6, 24 |
| Left Out of Social Activity | 902,669 | 801,505 | Subjective Well-Being, Loneliness, Life Satisfaction - Friends, Extraversion | 1, 18, 19, 23 |
| Life Satisfaction: Family | 906,592 | 289,950 | Life Satisfaction - Work, Subjective Well-Being, Life Satisfaction - Friends | 1, 23 |
| Life Satisfaction: Finance | 903,837 | 491,335 | Subjective Well-Being, Self-Rated Health, Loneliness | 1, 23 |
| Life Satisfaction: Friends | 903,837 | 401,682 | Life Satisfaction - Work, Subjective Well-Being, Left Out of Social Activity, Life Satisfaction - Family | 1, 23 |
| Life Satisfaction: Work | 906,592 | 702,208 | Subjective Well-Being, Life Satisfaction - Friends, Life Satisfaction - Family | 1, 23 |

| | | | | |
|---|---|---|---|---|
| Loneliness | 867,572 | 1,170,314 | Subjective Well-Being, Neuroticism, Left Out of Social Activity, Life Satisfaction - Finance, Depressive Symptoms | 1, 14, 18, 21, 23 |
| Neuroticism | 868,859 | 480,371 | Depressive Symptoms, Subjective Well-Being, Loneliness | 1, 14, 18, 21, 23 |
| Number Ever Born (Men) | 996,592 | 593,761 | Number Ever Born (Women), Age First Birth | 1, 6 |
| Number Ever Born (Women) | 996,592 | 497,396 | Number Ever Born (Men), Age First Birth | 1, 6 |
| Religious Attendance | 1,145,486 | 792,789 | Educational Attainment | 1, 5 |
| Risk Tolerance | 1,070,174 | 1,752,580 | Adventurousness | 1, 17 |
| Self-Rated Health | 1,134,460 | 1,250,433 | Life Satisfaction - Finance, COPD, Depressive Symptoms | 1, 14 |
| Self-Rated Math Ability | 1,135,947 | 663,024 | Highest Math, Cognitive Performance | 1, 4, 5 |
| Subjective Well-Being | 867,552 | 1,618,616 | Life Satisfaction - Work, Neuroticism, Loneliness, Left Out of Social Activity, Life Satisfaction - Friends, Life Satisfaction - Finance, Life Satisfaction - Family, Depressive Symptoms | 1, 14, 18, 21, 23 |

**Notes***:* "# SNPs" is the number of SNPs that were used to construct the PGI. "GWAS-equivalent *N*" is the GWAS-equivalent sample size as reported by MTAG. "Reference(s)" is the references for the GWAS summary statistics that were used to construct the PGI.

### 4. Self-rated health estimates

**Supplementary Table 4: Predictive accuracy of polygenic indices on self-rated health**

| **Polygenic Index** | **Beta** | **Standard Error** | ***P*-Value** |
|---|---|---|---|
| Externalizing single * | -0.11 | 0.02 | 1.13E-05 |
| Depressive Symptoms single * | -0.11 | 0.02 | 1.24E-05 |
| BMI single * | -0.10 | 0.02 | 1.48E-04 |
| ADHD single * | -0.10 | 0.02 | 1.22E-04 |
| Depression single * | -0.10 | 0.02 | 9.04E-05 |
| Number of children ever born women single * | -0.09 | 0.02 | 7.14E-04 |
| Type 2 diabetes single * | -0.09 | 0.02 | 2.67E-03 |
| Insomnia single * | -0.08 | 0.02 | 3.24E-03 |
| Neuroticism personality single * | -0.08 | 0.02 | 3.43E-03 |
| Cigarettes per day single * | -0.08 | 0.02 | 1.06E-02 |
| Systolic blood pressure single | -0.08 | 0.02 | 5.77E-02 |
| Ever smoke single * | -0.07 | 0.02 | 2.10E-02 |
| Left out of social activity single * | -0.07 | 0.02 | 2.81E-02 |
| Diastolic blood pressure single | -0.07 | 0.02 | 8.12E-02 |
| Triglycerides single | -0.05 | 0.02 | 6.10E-01 |
| Large artery stroke single | -0.05 | 0.02 | 2.05E+00 |
| Asthma single | -0.05 | 0.02 | 7.20E-01 |
| Migraine single | -0.04 | 0.02 | 2.87E+00 |
| Small vessel stroke single | -0.04 | 0.02 | 7.78E+00 |
| Any stroke single | -0.03 | 0.02 | 7.51E+00 |
| Any ischemic stroke single | -0.03 | 0.02 | 1.16E+01 |
| Narcissism personality single | -0.02 | 0.02 | 1.24E+01 |
| Asthma, Eczema, Rhinitis single | -0.02 | 0.02 | 1.60E+01 |
| Schizophrenia single | -0.02 | 0.03 | 2.33E+01 |
| Alcohol misuse single | -0.02 | 0.02 | 1.89E+01 |
| Extraversion personality single | -0.02 | 0.02 | 2.23E+01 |
| Alzheimers single | -0.01 | 0.02 | 2.82E+01 |
| Height single | -0.01 | 0.02 | 3.49E+01 |
| Atrial fibrillation single | -0.01 | 0.02 | 3.35E+01 |

#### 4. Self-rated health estimates

| | | | |
|---|---|---|---|
| Chronic kidney disease single | -0.01 | 0.02 | 3.72E+01 |
| Morning person single | -0.01 | 0.02 | 3.60E+01 |
| Breast cancer single | -0.01 | 0.02 | 3.74E+01 |
| Cannabis single | 0.00 | 0.02 | 4.67E+01 |
| Nearsightedness single | 0.00 | 0.02 | 5.14E+01 |
| Openess single | 0.00 | 0.02 | 4.89E+01 |
| Childhood reading single | 0.00 | 0.02 | 4.82E+01 |
| Cardioembolic stroke single | 0.00 | 0.02 | 4.79E+01 |
| Drinks per week single | 0.00 | 0.02 | 4.61E+01 |
| Risk tolerance single | 0.01 | 0.02 | 4.26E+01 |
| Hayfever single | 0.02 | 0.02 | 2.14E+01 |
| Friend satisfaction single | 0.02 | 0.02 | 1.50E+01 |
| Family satisfaction single | 0.03 | 0.02 | 7.61E+00 |
| Adventurousness personality single | 0.03 | 0.02 | 6.34E+00 |
| HDL single | 0.04 | 0.02 | 4.66E+00 |
| Age of first menses single | 0.05 | 0.02 | 1.19E+00 |
| Self-rated math ability single | 0.07 | 0.02 | 6.11E-02 |
| Physical activity single * | 0.07 | 0.02 | 3.10E-02 |
| Cognitive Performance single * | 0.07 | 0.02 | 3.09E-02 |
| Longevity single * | 0.09 | 0.02 | 4.02E-04 |
| Religious Attendance single * | 0.11 | 0.02 | 8.26E-06 |
| Highest math single * | 0.12 | 0.02 | 7.08E-07 |
| Subjective well-being single * | 0.13 | 0.02 | 3.82E-08 |
| Educational attainment single * | 0.14 | 0.02 | 4.78E-10 |
| Age of first birth single * | 0.14 | 0.02 | 6.57E-11 |
| Self-rated health single * | 0.20 | 0.02 | 1.04E-19 |

**Notes**: The coefficients in the table represent the estimated beta coefficients from each individual PGI regressed on self-rated health in the German Socioeconomic Panel Innovation Supplement. Self-rated health is the average self-rated health score, rated 1-5 with 1 being poor and 5 being very good, for each individual for all years they are observed in the survey. The observations are then collapsed into one observation per person and each PGI was

#### 4. Self-rated health estimates

[illegible] regressed separately given the high correlations between some PGIs. Overall, there were 2,229 individuals in each regression. Each regression included 5 year age bins for individuals 15 to 75 and a bin for individuals above age 75 interacted with an indicator for sex in addition to 20 genetic principal components. Starred PGI names indicate that the PGI is significant at a 5% significance level after a Bonferroni correction for 55 hypothesis. PGIs are sorted from largest to smallest in the table. For more information on how the PGIs are constructed, see table 3 in the main text.

Table 5. Observed height birth cohort effect estimates

| | **Beta** | **Standard Error** | ***P*-Value** |
|---|---|---|---|
| 1940's Birth Cohort | 0.06 | 0.62 | 9.25E-01 |
| 1950's Birth Cohort | 0.65 | 0.82 | 4.32E-01 |
| 1960's Birth Cohort | 2.151 | 0.99 | 3.02E-02 |
| 1970's Birth Cohort | 2.322 | 1.11 | 3.67E-02 |
| 1980's Birth Cohort | 3.090 | 1.43 | 3.03E-02 |
| Standardized height PGI | 3.08 | 0.14 | 1.33E-19 |
| Constant | 179.10 | 1.53 | 2.49E-40 |
| Observations | 6,465 | | |
| Number of individuals | 2,164 | | |
| R-squared | 0.614 | | |
| Sex FE | YES | | |
| Five Year Age Bin FE | YES | | |
| Interaction Sex and Age | YES | | |
| Clustered standard errors in Standard Error column | | | |

Notes: The coefficients in the table represent the estimated beta coefficients from observed phenotypic height in the German Socioeconomic Panel Innovation Supplement regressed on five year age bins, sex dummies, an interaction between sex dummies and the five year age bins, the standardized height PGI value, and birth cohort dummies. Standard errors are clustered for each individual, given that individuals are observed multiple times. Age bins are in five year intervals for individuals between the ages of 20 and 70. All individuals older than 70 are in one category. Individuals born before between 1923 and 1939 are all in the 1930s cohort, while individuals born after 1980 are all in the 1980 group. Individuals born between 1940-1949, 1950-1959, 1960-1969, and 1970-1979 are respectively labeled as 1940s, 1950s, 1960s, and 1970s. For more information on how the PGIs are constructed, see table 3 in the main text.

Table 6. Observed BMI birth cohort effect estimates

| | **Beta** | **Standard Error** | ***P*-Value** |
|---|---|---|---|
| 1940's Birth Cohort | -0.64 | 0.42 | 1.26E-01 |
| 1950's Birth Cohort | -1.70 | 0.54 | 1.78E-03 |
| 1960's Birth Cohort | -3.57 | 0.67 | 1.20E-07 |
| 1970's Birth Cohort | -3.76 | 0.82 | 4.59E-06 |
| 1980's Birth Cohort | -4.03 | 1.72 | 1.93E-02 |
| Standardized BMI PGI | 0.95 | 0.11 | 2.03E-09 |
| Constant | 26.67 | 1.76 | 2.55E-15 |
| Observations | 3,475 | | |
| Number of individuals | 1,458 | | |
| R-squared | 0.103 | | |
| Sex FE | YES | | |
| Five Year Age Bin FE | YES | | |
| Interaction Sex and Age | YES | | |
| Clustered standard errors in Standard Error column | | | |

Notes: The coefficients in the table represent the estimated beta coefficients from observed phenotypic BMI in the German Socioeconomic Panel Innovation Supplement regressed on five year age bins, sex dummies, an interaction between sex dummies and the five year age bins, the standardized BMI PGI value, and birth cohort dummies. Standard errors are clustered for each individual, given that individuals are observed multiple times. Age bins are in five year intervals for individuals between the ages of 20 and 70. All individuals older than 70 are in one category. Individuals born before between 1923 and 1939 are all in the 1930s cohort, while individuals born after 1980 are all in the 1980 group. Individuals born between 1940-1949, 1950-1959, 1960-1969, and 1970-1979 are respectively labeled as 1940s, 1950s, 1960s, and 1970s. For more information on how the PGIs are constructed, see table 3 in the main text.
